# SenNet Portal: Build, Optimization and Usage

**DOI:** 10.64898/2026.02.06.704469

**Authors:** Katy Börner, Philip D. Blood, Jonathan C. Silverstein, Matthew Ruffalo, Rahul Satija, Nils Gehlenborg, Brendan Honick, Andreas Bueckle, Yashvardhan Jain, Danial Qaurooni, Bill Shirey, Max Sibilla, Kay Métis, Joe Bisciotti, Richard S. Morgan, David Betancur, Ryan Sablosky, Morgan L. Turner, So-Jin Kim, Patty J. Lee, Josh Bartz, Sergii Domanskyi, Samuel T. Peters, Archibald Enninful, Negin Farzad, Rong Fan, SenNet Team, Bruce W. Herr

**Affiliations:** Department of Intelligent Systems Engineering, Indiana University, Bloomington, IN, USA; CIFAR MacMillan Multiscale Human program, CIFAR, Toronto,ON, Canada; Pittsburgh Supercomputing Center, Carnegie Mellon University, Pittsburgh, PA, USA; Department of Biomedical Informatics, University of Pittsburgh School of Medicine, Pittsburgh, PA, USA; Ray and Stephanie Lane Computational Biology Department, Carnegie Mellon University, Pittsburgh, PA, USA; New York Genome Center, New York, NY, USA; Department of Biomedical Informatics, Harvard Medical School, Boston, MA, USA; Division of Pulmonary, Critical Care, and Sleep Medicine, Department of Medicine, Icahn School of Medicine at Mount Sinai, New York, NY, USA; Masonic Institute on the Biology of Aging and Metabolism, Department of Genetics, Cell Biology, and Development, University of Minnesota, Minneapolis, MN, USA; The Jackson Laboratory for Genomic Medicine, Farmington, CT, USA; Masonic Institute on the Biology of Aging and Metabolism, Department of Biochemistry, Molecular Biology, and Biophysics, University of Minnesota, Minneapolis, MN, USA; Department of Biomedical Engineering, Yale University, New Haven, CT, USA; Department of Pathology, Yale University School of Medicine, New Haven, CT, USA

**Author notes:** Corresponding authors: Katy Börner, Philip D. Blood, Jonathan C. Silverstein, Bruce W. Herr II. Full list of authors included in SenNet Team is available at the end of the paper.

## Abstract

Cellular senescence is a hallmark of aging and a driver of functional decline across tissues, yet its heterogeneity and context dependence have limited systematic study. The Common Fund’s Cellular Senescence Network (SenNet) Program addresses this challenge by generating multimodal, multi-tissue datasets that profile senescent cells across the human lifespan and complementary mouse models. The SenNet Data Portal (https://data.sennetconsortium.org) serves as the public gateway to these resources, providing open access to harmonized single-cell, spatial, imaging, transcriptomic, and proteomic data; senescence biomarker catalogs; and standardized protocols that can be used to comprehensively identify and characterize senescent cells in mouse and human tissue. As of April 2026, the portal hosts 2,041 publicly available human and mouse datasets across 15 organs using 6 general assay types. Experts from 13 Tissue Mapping Centers (TMCs) and 12 Technology Development and Application (TDAs) components contribute tissue data, analyze data, identify senescent biomarkers, and agree on panels for cross-tissue antibody harmonization. They also register human tissue data into the Human Reference Atlas (HRA) and develop user interfaces for the multiscale and multimodal exploration of this data. Built on a scalable hybrid cloud microservices architecture by the Consortium Organization and Data Coordinating Center (CODCC), the Portal enables data submission, management, integrated analysis, spatial context mapping, and harmonized access to cross-species data critical for aging research. This paper presents user needs, the Portal’s architecture, data processing workflows, and senescence-focused analytical tools; usage scenarios illustrating applications in biomarker discovery, quality benchmarking, hypothesis generation, spatial analysis, cost-efficient profiling, and cell distance distribution analysis; and utility and usage by the larger researcher community. Current limitations and planned extensions—including expanded spatial-omics releases and improved tools for senotype characterization—are discussed. SenNet protocols, code, and user interfaces are freely available on https://docs.sennetconsortium.org/apis.

## Main

Inaugurated in 2021, the Common Fund’s Cellular Senescence Network (SenNet) Program was established to comprehensively identify and characterize senescent cell identities and abundances across the body, across various states of human health, and across the lifespan^1,2^. Resources and results currently being produced by SenNet include publicly accessible atlases of senescent cells and descriptions of their shared and distinct molecular identities, see https://sennetconsortium.org for resources, data, and publication menus. Data collected from human and mouse model organism tissues were used to identify and characterize senescent cells.^3^ Novel tools and technologies were designed, implemented, and validated in close collaboration with experts from while expanding methods and technologies developed by the Common Fund’s Human Biomolecular Atlas Program^4,5^ and Single Cell Analysis Program.^6^

There are three SenNet portal websites: The SenNet Consortium Website (https://sennetconsortium.org) introduces the SenNet consortium and provides public access to resources, news, working groups, and consortium member services. The SenNet Data Portal (https://data.sennetconsortium.org) makes experimental data easy to access and download, modeled after the HuBMAP Data Portal^7^. The cross-consortium Human Reference Atlas (HRA) Portal (https://humanatlas.io) makes reference atlas data and code freely available in support of data harmonization, mapping, and multiscale data exploration. All data are shared according to FAIR principles (Findable, Accessible, Interoperable, and Reusable) to maximize data utility.^8^ The HRA and SenNet data portals are synergistic: a reference atlas cannot be constructed without access to high quality experimental data and the atlas is used to harmonize and federate experimental data (e.g., spatially register tissue in a 3D common coordinate framework; to design and use standardized and validated antibody panels; to update ontologies to include novel cell types) so it can be compared across teams, organs, or assay types.

SenNet united cellular senescence researchers within the program and beyond to agree on key terminology, see **Box 1**. Additional terms are defined in the Getting Started with Data Submission guide^9^ and the HRA Glossary.^10^

#### Box 1. Key SenNet Terminology

- **Common coordinate framework (CCF):** Consists of ontologies and reference object libraries, computer software (e.g., user interfaces) and training materials that (1) enable biomedical experts to semantically annotate tissue samples and to precisely describe their locations in the human/mouse body (“registration”), (2) align multi-modal tissue data extracted from different individuals to a reference coordinate system (“mapping”), and to (3) provide tools for searching and browsing experimental data at multiple levels from the whole body down to single cells (“exploration”).^11^
- **Data curators:** Curators work with a funded component’s designated data submitter to ingest data according to SenNet standards.
- **Dataset:** The files and metadata that comprise and describe a run of an experimental assay. Primary datasets contain the raw data and primary processing while derived datasets contain additional assays output from processing primary datasets (e.g., for spatial assays, there may be histology and -omics derived datasets).
- **Data submitters:** Submitters work closely with curators to upload experimental data via the SenNet Ingest Portal.
- **Data upload:** One or more datasets generated by the same primary assay type submitted together to the Consortium Organization and Data Coordinating Center (CODCC).
- **Metadata:** ‘Data about data’ refers to descriptions about sources, tissue samples, and datasets. Metadata for a particular purpose can be defined by a metadata schema. SenNet and HuBMAP co-develop and use the same schemas (except for an additional murine metadata schema for SenNet), and modifications can be made by either consortia’s Data Coordination Working Groups.
- **Provenance:** The origins of each entity, captured internally via a graph database. A typical flow of provenance in SenNet is source -> organ -> tissue block -> a further block, section, or suspension -> dataset.^12^ Everything from organ down is registered to a direct ancestor (organ registered to source (individual human or murine organism), tissue block to organ, section/suspension/block to first block, dataset to sample). A dataset is descended from the most granular sample against which the experimental modality was run.
- **Reference atlas:** A comprehensive, high-resolution, three-dimensional, multiscale atlas of all the cells in the healthy human/mouse body. The Human Reference Atlas (HRA) provides standard terminologies and data structures for describing specimens, biological structures, and spatial positions linked to existing ontologies.^13^
- **Senescence-Associated Secretory Phenotype (SASP):** The secretion by senescent cells of pro-inflammatory cytokines, growth factors, chemokines, proteases, and other bioactive molecules^3^. Phenotypic expressions of senescent cells that reshape tissue microenvironments and influence nearby cells.
- **Senescent cells:** Cells that have undergone a stable, near-permanent cell-cycle arrest, remain metabolically active, resist apoptosis, and elaborate a pro-inflammatory secretome (SASP) and/or other phenotypic changes that contribute to aging and age-related disease^3^.
- **Secondary senescence:** Senescence transmitted to neighboring cells via either SASP or direct cell-to-cell signaling^3^.
- **Senescence microenvironment (SME):** Anatomical sites and cellular neighborhoods enriched in senescent cells and their SASP effects, including the reshaping of tissue architecture, the remodeling of the extracellular matrix, and the modulation of immune surveillance^2^.
- **Senolytics:** Senotherapeutic drugs that selectively kill senescent cells^14,15^.
- **Senotype:** A senescent-cell subtype with a distinct, integrative molecular and functional profile, shaped by cell lineage, tissue microenvironment, inducing stimulus, and time since senescence onset^2^.
- **Subclinical senescence**: The early and potentially invisible stages of cellular aging that involve histologically normal cells exhibiting subtle senescence signatures that may be detectable through high-resolution assays.
- **Validation**: The process by which curators check a data upload’s metadata (according to a metadata schema), directory and file structure (according to a directory schema) for compliance with CODCC standards.

## Results

As of January 2026, SenNet results comprise (1) a set of well-defined user needs; (2) a scalable hybrid cloud microservices architecture; (3) data obtained through well-defined data submission and ingestion procedures; (4) generalized data processing and analysis pipelines; (5) reference atlas construction and usage workflows; (6) senescence focused data visualizations and user interfaces; and (7) documentation and demonstration resources. All seven contributions are detailed subsequently together with five SenNet usage scenarios that demonstrate how SenNet data and tools can be used to advance basic research and clinical practice.

### General User Needs

To better understand how the SenNet Data Portal supports real-world research, we approached clinicians, experimental and computational biologists whose labs had contributed data to the Portal. In ten semi-structured interviews, we discussed existing and potential user goals, workflows, and challenges. Interview transcripts were analyzed using an inductive approach to discover recurring use cases, which were then developed into detailed user stories in collaboration with the data contributors. The analysis generated seven representative user stories reflecting key use cases across three domains of discovery, reference, and translation, see **Table 1**. Each story outlines one of the Portal’s value propositions and illustrates the diverse ways users could engage with SenNet resources in their work. In four cases, we followed these interviews with more in-depth discussions focused on understanding Portal workflows in more depth. These conversations generated context-rich, illustrated *Usage Scenarios* which are presented in the ***Results*** section.

**Table 1:** User stories. Feature summary, target user roles, user activities, and added value for seven user stories that are significant for SenNet Portal users.

| Area | User Role | User Activity | SenNet Portal Added Value |
|---|---|---|---|
| Discovery Use Cases |  |  |  |
| **US#1.** Exploratory Analysis | Researchers, Clinicians | Analyzing preliminary findings about gene expression and tissue architecture | View and compare gene expression and tissue structure across tissues, age groups, and species to better interpret early findings |
| **US#2.** Integrative Analysis | Researchers, Clinicians | Investigating how circulating molecular signals influence dysfunction across organs | Support systems-level research by integrating senescent cell profiles and circulating signals across tissues, enabling discovery of cross-organ mechanisms of dysfunction |
| Reference Use Cases |  |  |  |
| **US#3.** Data Validation | Researchers | Identifying senescence in particular tissues using specific markers | Use reference datasets to validate findings in unfamiliar tissues or conditions, reducing the need to generate new data |
| **US#4.** Normal Tissue Comparison | Pathologists | Examining gene expression and tissue architecture to discover early disease markers | Compare samples to histologically normal reference tissues to detect early markers of disease in at-risk populations |
| **US#5.** Tool Evaluation | Computational Researchers | Evaluating the performance of existing machine learning models to develop benchmarking tools for senescence | Use diverse, multi-tissue datasets to train and evaluate machine learning models, improving accuracy and reproducibility in senescence detection |
| Translational Use Cases |  |  |  |
| **US#6.** Therapy Development | Researchers, Clinicians | Identifying shared and tissue-specific pathways of aging | Access harmonized human and mouse senescence datasets under shared ontologies and visually compare expression within species to surface candidate pathways for therapeutic targeting |
| **US#7.** Cross-Species Translation | Researchers, Clinicians | Translating well-characterized signaling pathways from mouse to human | Retrieve co-located mouse and human datasets of matched organs and assays under common metadata schemas, enhancing the relevance of preclinical models |

### Flexible hybrid cloud microservices architecture

The SenNet Consortium Organization and Data Coordination Center (CODCC) developed a flexible hybrid cloud microservices architecture^4,5^ based on the HuBMAP Consortium^4,5^ and data portal to support data curation, ingestion, integration, access, analysis, exploration, and download (https://portal.hubmapconsortium.org). See cloud microservices architecture details in a related HRA publication^16^ and flow of data through Portal architecture from data submission to consumption in the HuBMAP Portal paper. This microservices architecture was adopted for the SenNet Portal (https://data.sennetconsortium.org) on distinct infrastructure, demonstrating the approach is transferable among supercomputer centers, public cloud, university infrastructure, and mixed/hybrid deployments. Data are ingested using the *Ingest UI* (**Fig. 1**, e.g., the Ingest UI is made up of parts of the Portal Application, Ingest API, and Ingest Pipeline, while leveraging other APIs). The HRA Registration User Interface (*RUI*)^11,17,18^ supports tissue registration in 3D so tissue blocks can be explored within the 3D human reference body using the Exploration User Interface (*EUI*)^11^. Data are then processed and analyzed using diverse cell segmentation and cell type annotation tools, including *Azimuth*^19^ and made available via the Portal. *Vitessce*^20^ supports the exploration of multi-modal spatial and single cell data^7^, see example in usage scenario section ***Hypothesis Generation***.

**Figure 1:**
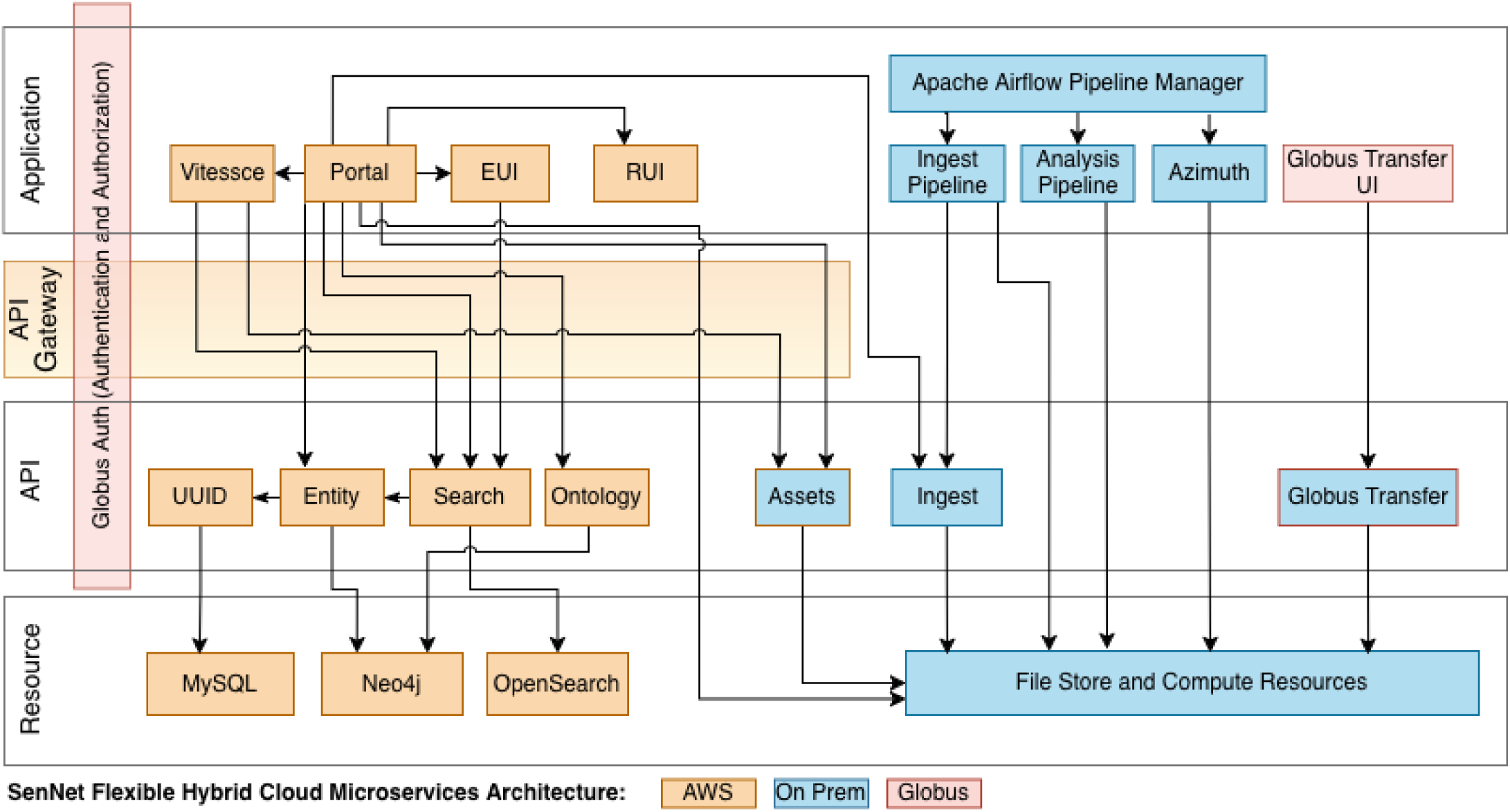
SenNet infrastructure architecture and functionality. On premises resources at the University of Pittsburgh and public cloud resources in AWS are used to store, ingest, process and expose the data publicly via the SenNet Data Portal. APIs exist for all programmatic discovery and access to datasets. The API Gateway monitors all API access and allows only authorized access to SenNet provenance and metadata. Applications support data ingest, analysis, visualization, search, and download via diverse user interfaces. Globus Auth is used for all authentication and authorization to non-public SenNet APIs, applications, and data. All data are available for download via Globus Collections. Data upload and download are managed via Globus Transfer. EUI: Exploration User Interface^11^ ; RUI: Registration User Interface^11^; UUID: universally unique identifier; On Prem: on premises.

Major advances of SenNet beyond the HuBMAP data portal include: The SenNet Data Portal integrates ingest functionality into the public data portal (HuBMAP has separate Ingest and Data portals). RUI-based tissue registration is required for all human SenNet data (in HuBMAP, 3D block registration is encouraged). SenNet supports mouse and human sources, including senescence-induced and diseased (HuBMAP contains healthy human tissue). SenNet supports additional tissues (e.g., muscle, adipose, mammary gland) and improved navigation and visualization of provenance information. Provenance and data flow visualizations are directly interactive for investigative feedback, orientation to data and provenance, and search in SenNet, using the framework of data-driven documents, substantially enhancing user experience.

### Data Submission, Ingestion, and Access

Data submission and ingestion in SenNet is curated by the CODCC. Submission begins with a bulk data upload by a data submitter. Ingestion is the process of the curator handling data submitters’ bulk data uploads via validation, reorganization, central analysis, quality assurance, and publication of the data and metadata. SenNet adopted HuBMAP’s workflows, metadata and directory schema^21^, and co-developed new schemas. SenNet has also developed a unique mouse source metadata schema^22^ since it supports both human and mouse tissue samples.

Datasets submitted to the CODCC within bulk uploads require at least two metadata files, one describing the assay and the other recording applicable contributor information. Data are organized according to the directory and file structure described in the relevant directory schema for the assay. Preparing an upload through data publication involves collaboration among curators and data submitters. Avenues for collaboration include weekly SenNet Data Coordination Working Group (DCWG) meetings to set standards and share pragmatic approaches throughout the process, twice-a-week curator office hours, and triage in cooperation with the SenNet Help Desk. Curators and submitters work together to publish experimental protocols on SenNet’s protocols.io workspace^23^ (218 distinct protocols to date), register entities in the provenance chain (sources, organs, and tissue samples), prepare and ingest metadata, and submit spatial coordinate information for human tissue blocks registered in the HRA’s Registration User Interface.^11^

In April, 2026, SenNet generated and published 1,412 human and 629 mouse primary datasets for a total of 2,041 datasets using microscopy (810 datasets), antibody-based imaging (81), mass spectrometry (179), transcriptomics (587), spatial transcriptomics (615), and epigenomics (108) technologies. These include 175 different tissues (∼40% mouse and 60% human). The SenNet data portal provides an interactive diagram of these data sets that link to the primary data ingested (**Fig. 2**).

**Figure 2.**
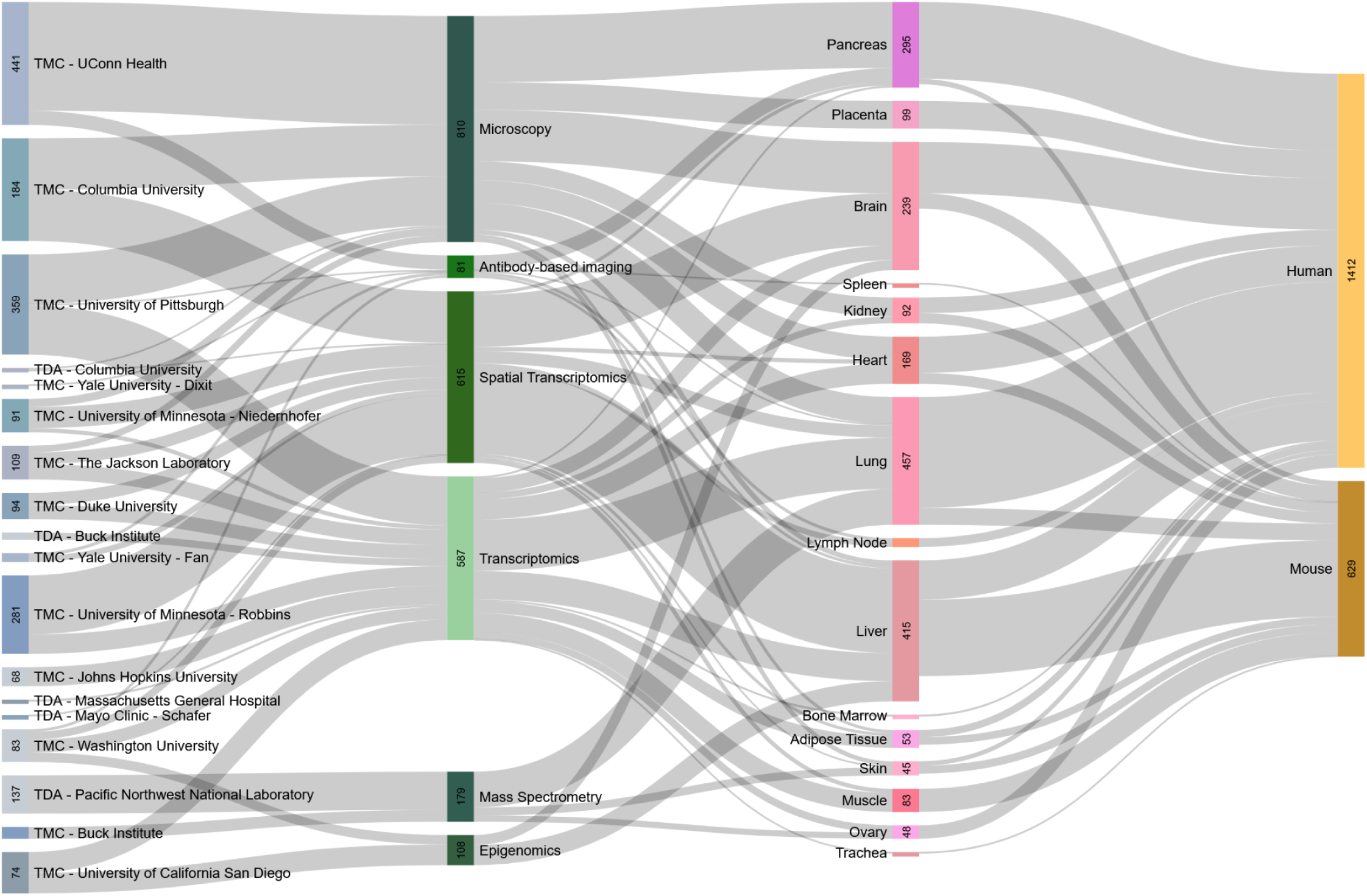
Sankey diagram of the SenNet dataset generation. The diagram illustrates the flow of datasets from data upload to publication. The vertical size of each colored bar and flow (grey bands) shows the proportion of datasets for each source, technology, tissue and species. From left to right, 17 Tissue Mapping Centers (TMCs) and Technology Development and Application Projects (TDAs) are submitting data for 6 general assay types (microscopy, antibody-based imaging, mass spectrometry, transcriptomics, spatial transcriptomics, and epigenomics), for 175 tissue types in humans or mice. An interactive version of the Sankey graph with direct links to data is available at https://docs.sennetconsortium.org/data-sankey.

**Figure 3.**
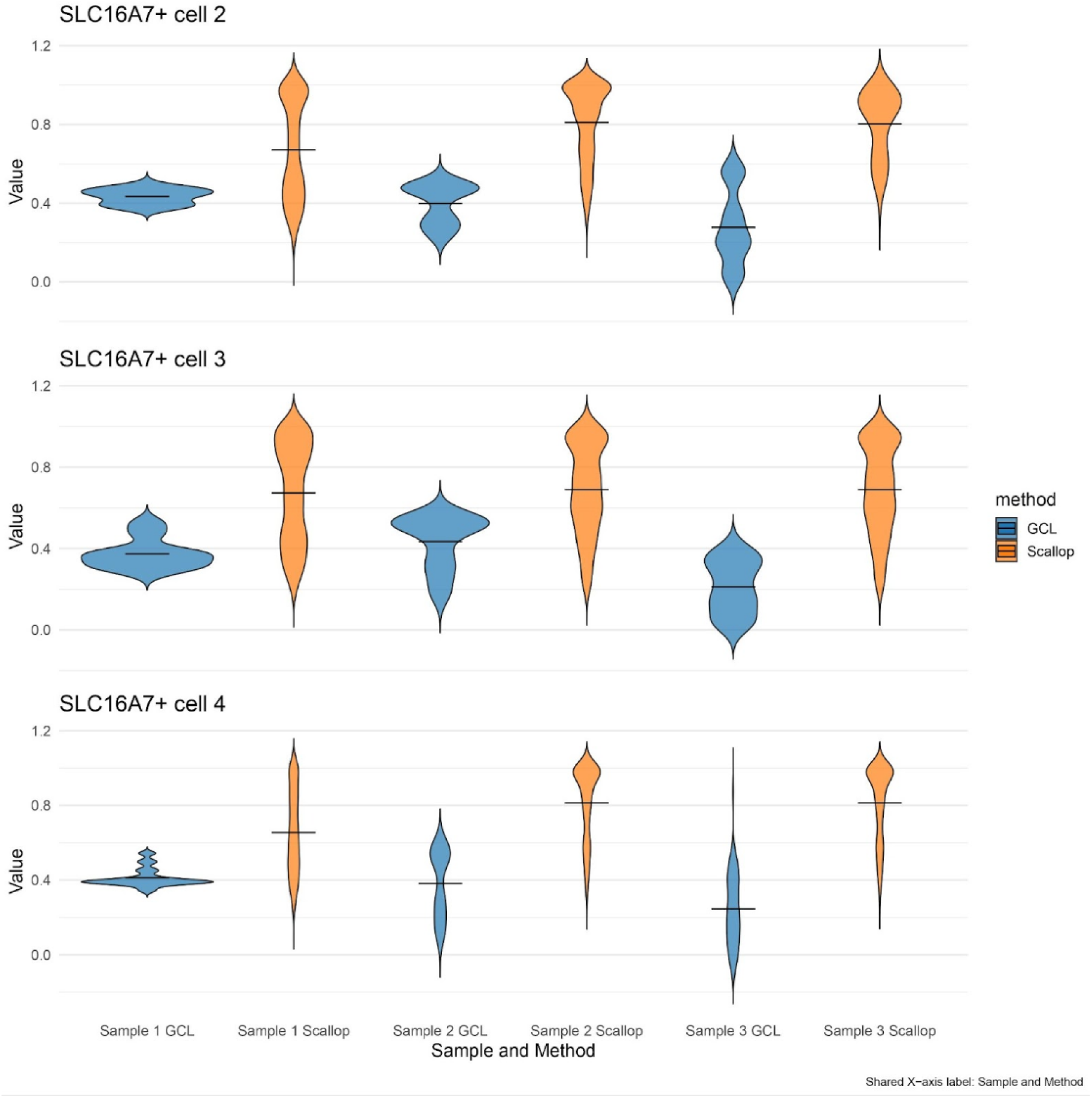
Benchmark comparisons of Global Coordination Level and Scallop. The high-quality lung transcriptomic dataset provided by SenNet facilitates robust benchmarking of algorithms, including those designed to quantify transcriptional noise, across a broad range of cell types. Processed data from following SenNet datasets were used in this example: SNT246.PGHH.774, SNT449.GRWG.598, and SNT397.SPGW.436.

### Data Processing and Analysis

The CODCC includes focus on data ingestion, analysis, and visualization workflow integration and tool development, development of Azimuth references^24^, and the Human Reference Atlas^25^ in close collaboration with external experts. The SenNet portal hosts datasets processed through uniform computational pipelines, one per assay type. This includes a single-cell/single-nucleus transcriptomic quantification pipeline built on the Salmon method^26^ and ScanPy for downstream analysis^27^, single-cell quantification of open chromatin regions (scATAC-seq) built on ArchR^28^, and cell and nucleus segmentation of multichannel images performed by the top-performing segmentation method for each particular assay^29^, then unsupervised downstream analysis via the SPRM package^30^. When possible, uniform processing pipelines include senescence-specific analysis, such as the inclusion of DeepScence^31^ into the sc/snRNA-seq pipeline, producing predictions of cellular senescence for all sc/snRNA-seq datasets that have been processed since the inclusion of this pipeline step. This uniform processing of datasets between providers, assay variants, and tissues allows for easier integrative analysis, removing artificial barriers to data harmonization, and supports the senescence analysis detailed in other sections of this paper.

### SenNet Atlas Construction and Usage

Spatially registered and ontology-aligned SenNet data are used to construct a reference atlas in support of senescence studies, see **Box 1** for terminology. In April 2026, there exist 2,306 spatially registered human datasets for 1,018 tissue sections, cut from 679 tissue blocks with 479 extraction sites in 12 organs from 294 donors, uploaded by 10 tissue data providers, see Exploration User Interface (EUI, https://apps.humanatlas.io/eui)^11^ with SenNet filter applied. This all-HRA EUI does include secondary datasets, (e.g., cell by gene matrices or cell type annotation and segmentation masks) computed from the primary data; and it uses data from the HRA KG which is updated every six months with the HRA release. The all-HRA EUI provides access to a total of 9,067 tissue datasets, with 2,306 (or 25.4 percent) coming from SenNet.

SenNet’s EUI at https://data.sennetconsortium.org/ccf-eui only includes primary human datasets, analogous to counts on the SenNet Data Portal. It supports easy access to 1,307 public datasets from 290 donors. With consortium login, 1,692 datasets from 507 donors are accessible and will become available publicly after data validation has been completed.

Exactly 54 of these datasets were used to compute HRA cell type populations (HRApop) v1.0 for anatomical structures^32^ in June 2025 (52 from the lung and two from the liver). The other 988 datasets were either not in existence when HRApop v1.0 was computed or came from organs not covered by existing cell type annotation tools (e.g., 86 datasets from placenta and 65 datasets from the ovaries). That is, 8.1% of the HRApop v1.0 data come from SenNet, making it a major data source for HRApop construction. Using HRApop, cell type populations and biomarker expression values can be predicted for HRA-registered 3D tissue volumes—before any single cell assay type is run; inversely, given a cell type by biomarker expression matrix, the 3D volume from which this tissue was cut can be predicted.

Five CODEX human lymph node datasets from SenNet with 8,918,854 cells were used in a recent cross-consortium study of spatially resolved omics data that explores endothelial cell environments across 14 studies comprising 12 tissue types and a total of 47,349,496 cells^33^ All data and code are freely available and the Cell Distance Explorer (CDE, https://apps.humanatlas.io/cde) has been used to analyze other SenNet human and mouse datasets.

### Data Visualizations and User Interfaces

Different user interfaces exist to explore experimental tissue datasets and reference atlas data.

#### User interfaces to experimental data

##### Vitessce

Visualization Tool for Exploration of Spatial Single-Cell Experimentation (Vitessce) is an open-source, web-based framework designed for the integrative visualization of multi-modal, spatially resolved single-cell data in both 2D and 3D (https://vitessce.io). It enables researchers to interactively explore high-resolution imaging supported by Viv^34^, spatial transcriptomics, and conventional single-cell data in a linked visualization interface. A diverse set of Vitessce visualizations, spanning spatial and non-spatial as well as single- and multi-modal data, was originally developed for the HuBMAP Data Portal to support specific assay requirements and analytes across 16 data types. These visualizations are also fully supported by the SenNet Data Portal, see the ***Hypothesis Generation*** section for exemplary Vitessce visualizations used with SenNet data.

#### User interfaces to reference atlas data

There are diverse user interfaces (UIs) that support atlas construction (e.g., authoring ASCT+B tables or OMAPs; spatial and semantic registration of tissue blocks) and UIs that support atlas usage (e.g., predicting cell type populations for registered tissue blocks). We explain the former first, realizing that few experts will construct the HRA, most experts will use the HRA.

##### ASCT+B Reporter with Organ Mapping Antibody Panel (OMAP) Support

The ASCT+B Reporter supports the visualization of ASCT+B tables^35^ in the form of trees showing anatomical structures (AS) in the human body, the cell types (CT) located in them, and the biomarkers (B) used to characterize these cell types (apps.humanatlas.io/asctb-reporter). A mode was added to explore, compare, and optimize OMAPs^36^ specific to organs, multiplexed antibody-based imaging methods, and tissue preservation methods. The ASCT+B Reporter is critical for authoring, reviewing, and optimizing ASCT+B tables and OMAPs.

##### SenNet-branded Registration User Interface (RUI)^11^

The SenNet Data Portal (data.sennetconsortium.org) enables data providers to register data sources (donors), organs, tissue blocks, etc. A RUI instance, embedded as a web component, is available during this registration workflow to indicate the spatial origin (also called extraction site) of a tissue block inside one of the 73 3D reference objects in HRA v2.3 representing the organs of the healthy, adult human body. Details are available in the data submission guide.^37^

##### SenNet-branded Exploration User Interface (EUI^11^)

Enables the spatial and semantic exploration of SenNet donors, tissue blocks, tissue sections, and datasets if they have been annotated with an extraction site via the RUI. Each extraction site has a 3D position, rotation, and size. The SenNet specific EUI is available at data.sennetconsortium.org/ccf-eui, the all-HRA EUI is at apps.humanatlas.io/eui. See example in usage scenario section ***Cell Distance Distributions***.

##### Cell Distance Explorer (CDE^33^)

Enables a user to visualize cell-to-nearest-anchor-cell linkages (apps.humanatlas.io/cde) and distance distributions. The user provides a CSV file with a list of cells (x, y, z-position, plus a cell type label) and metadata (e.g., visualization title, imaging technology, organ, sex) and designates an anchor cell type (e.g., endothelial cells). The CDE then computes links between each cell type and its nearest anchor cell. As auxiliary visualizations, the CDE provides a histogram of distances, color-coded by cell type (and a bucket for all), as well as one violin plot per cell type with an inline box-and-whisker plot. The CDE allows researchers to analyze and compare changes in cell-cell distance distributions across tissue regions, between normal and diseased tissue, and across diseases. It thus enables exploration of how cell organization is impacted by different conditions. See example in usage scenario section ***Cell Distance Distributions***.

##### Cell Population Graphs (apps.humanatlas.io/cell-population-graphs)

Make it possible to compare cell type populations across experimental datasets and in relation to HRApop atlas datasets. Each stacked bar graph presents an experimental or atlas dataset; different cell types are rendered in different colors and bar height corresponds to cell count or percentage. Bars can be sorted by total cell count or prevalence of specific cell types, and they can be grouped by dataset-specific properties (e.g., sex, developmental stage, ethnicity if metadata is available). A newer interface is the HRApop Visualizer (apps.humanatlas.io/hra-pop-visualizer), which enables exploration of cell type populations for anatomical structures, extraction sites, and datasets in HRApop v1.0.^32^

##### HRA Organ Gallery virtual reality (VR)

A stand-alone VR application (http://humanatlas.io/hra-organ-gallery), available free of charge in the Meta Store.^38^ Users can view 77 3D reference objects and 2,894 tissue blocks linked to 9,067 datasets from 902 donors across 20 consortia—including 679 tissue blocks from SenNet. Additionally, two SenNet datasets (CODEX, Visium) can be explored in immersive 3D visualizations.

### Documentation and Demonstrations

The SenNet portal (https://sennetconsortium.org) provides five main dimensions of engagement: **resources**, **data exploration**, **news**, **involvement**, and **member services**.

The ***Resources*** section is the technical backbone for contributing and using SenNet data. It enables users to access biomarker lists, documentation of software tools, the Human Reference Atlas, experimental protocols on protocols.io, SenNet publications in PubMed and via Google Scholar profile with citations, software on GitHub (https://github.com/sennetconsortium), as well as summaries of novel technologies and tools developed within SenNet.

The **Data** section, which constitutes the core of the portal, is organized around submitting and accessing datasets. It supports contributor workflows by offering clearly defined entry points for registering specimens, uploading data, and validating metadata through standardized templates. The portal provides detailed documentation for these processes and enables contributors to monitor submission status to ensure compliance with consortium-wide quality controls. (Submitted datasets are indexed under a shared metadata schema and aligned to common anatomical and cell type ontologies.) The Data section also provides access to curated and published datasets. The portal offers three entry points for new users. They can familiarize themselves using onboarding materials, explore interactive data previews through the dynamic SenNet Sankey diagram (https://data.sennetconsortium.org), or explore faceted search organized by popular categories. More experienced users can search and filter datasets by entity type (sample, dataset, source, etc.), tissue, assay, contributor, or organ, view integrated summaries and visualizations, and export results for downstream analysis. In addition to web-based interfaces, the portal supports programmatic access through Application Programming Interfaces (APIs) at https://entity.api.sennetconsortium.org. The ***Usage Scenarios*** section illustrates how these tools can be combined for data exploration, integration, and reuse across diverse research contexts.

**News** chronicles the consortium’s ongoing activities, including both internal milestones and public-facing achievements. Common news items include announcements, events, funding opportunities, and new publications^39^. A subsection of News, the portal’s Interview Series, features conversations with scientists and project leaders that trace how different efforts within SenNet connect and progress over time (https://sennetconsortium.org/interview-series). Another subsection, the SenNet Sentinel, is dedicated to a monthly newsletter that spotlights announcements, publications, programs, and personnel for internal and external audiences.

The ***Involvement*** section invites researchers and institutions to take part in SenNet’s collective work in one of two ways. They can either apply to become Associate Members of the consortium (of which there are eight in January 2026), where they would contribute to its research goals directly. Alternatively, users can join working groups dedicated to benchmarking, data coordination, atlas construction, or emerging technologies, allowing them to participate in shaping the methods and standards that guide SenNet’s evolving research practices.

Finally, the **Member services** section forms the administrative core of the Portal and handles internal communication and coordination between consortium members. Consortium members access their user profiles through a secure log-in, upload and review experimental data for publication, follow internal calendars, and share meeting materials and documents.

### Usage Scenarios

General user needs were detailed in the ***General User Needs*** section. Here, we present five exemplary data and tool usage scenarios that showcase how SenNet data can be used to (1) benchmark and improve noise cleaning algorithms and tools, (2) showcase that pilot transcriptomics lung data in mouse is supported by standardized, consortium-generated datasets, (3) guide pathologists in spatial and other data analyses in diabetes in human pancreas with a focus *Islets of Langerhans* functional tissue units (FTUs), (4) run a temporal metastudy, and (5) compute cell distance distributions for immune cells. These scenarios were derived via extended interviews with data contributors to the Portal. They are presented as illustrative examples of how the Portal can serve diverse use cases.

#### Data Quality Benchmarking

A long-standing challenge for senescence researchers is that cellular senescence is not defined by a single biomarker, but by multiple biomarkers and signatures that are highly specific to cell type and (see subclinical senescence in **Box 1**). This diversity results in a weak signal for detecting senescence that is difficult to differentiate from other signals or technical noise. The problem is compounded once we consider the strong batch effects that characterize transcriptomic datasets compiled across multiple studies and originating from differences in protocols, lab environments, or sequencing technologies.^40^ Researchers have often had to resort to crude normalization methods that risk erasing precisely the subtle differences in biological signals they hope to study, including variations in senotypes. SenNet’s carefully curated data, generated in a coordinated manner across a small set of labs using standardized protocols, offers something rare: *large-scale, high-quality transcriptomic data with low technical noise*. For a meta-study involving multiple organs, Bartz and colleagues tapped into SenNet’s harmonized raw datasets, using them both to benchmark existing noise-measurement algorithms and to train data-hungry deep learning models capable of distinguishing emerging senescence micro-environments. They benchmark methods such as Global Coordination Level Analysis (GCL)^41,42^ and Scallop^43^ for transcriptional noise, alongside senescence detection tools like hUSI^44^, SENCAN^45^, and SenCID.^46^

#### Hypothesis Generation

A group of pulmonary biomedical researchers turn to the SenNet portal to strengthen their resubmission of a major grant which focuses on senescence markers in lung tissue. The portal helps Drs. Lee and Kim in two major ways. First, its standardized datasets and user-friendly interface allow them to stress-test their hypothesis. Their menu of options includes using UMAP visualization and gene expression panels to test candidate genes and identify cell clusters with high expression levels (**Fig. 4a**), comparing expression levels of senescence markers between custom groups (**Fig. 4b**), along with other comparisons based on age, sex, and modality. They can also incorporate other datasets from the same upload batch to probe whether candidate markers participate in tissue-specific aging pathways or reflect broader aging signatures across cell types. Evaluating their preliminary findings against benchmark datasets empowers these researchers to confirm, refine, or revise their initial hypotheses. The portal also supports the formulation of new hypotheses. While looking for other senescence-related markers and comparing expression levels between user-defined groups, users might observe enriched expression within certain subpopulations, suggesting a potential functional synergy between these two molecules in the mouse lung. This data-driven insight motivates them to consider novel hypotheses, such as whether concurrent blockade of these targets can offer synergistic therapeutic benefits in lung injury models.

**Figure 4.**
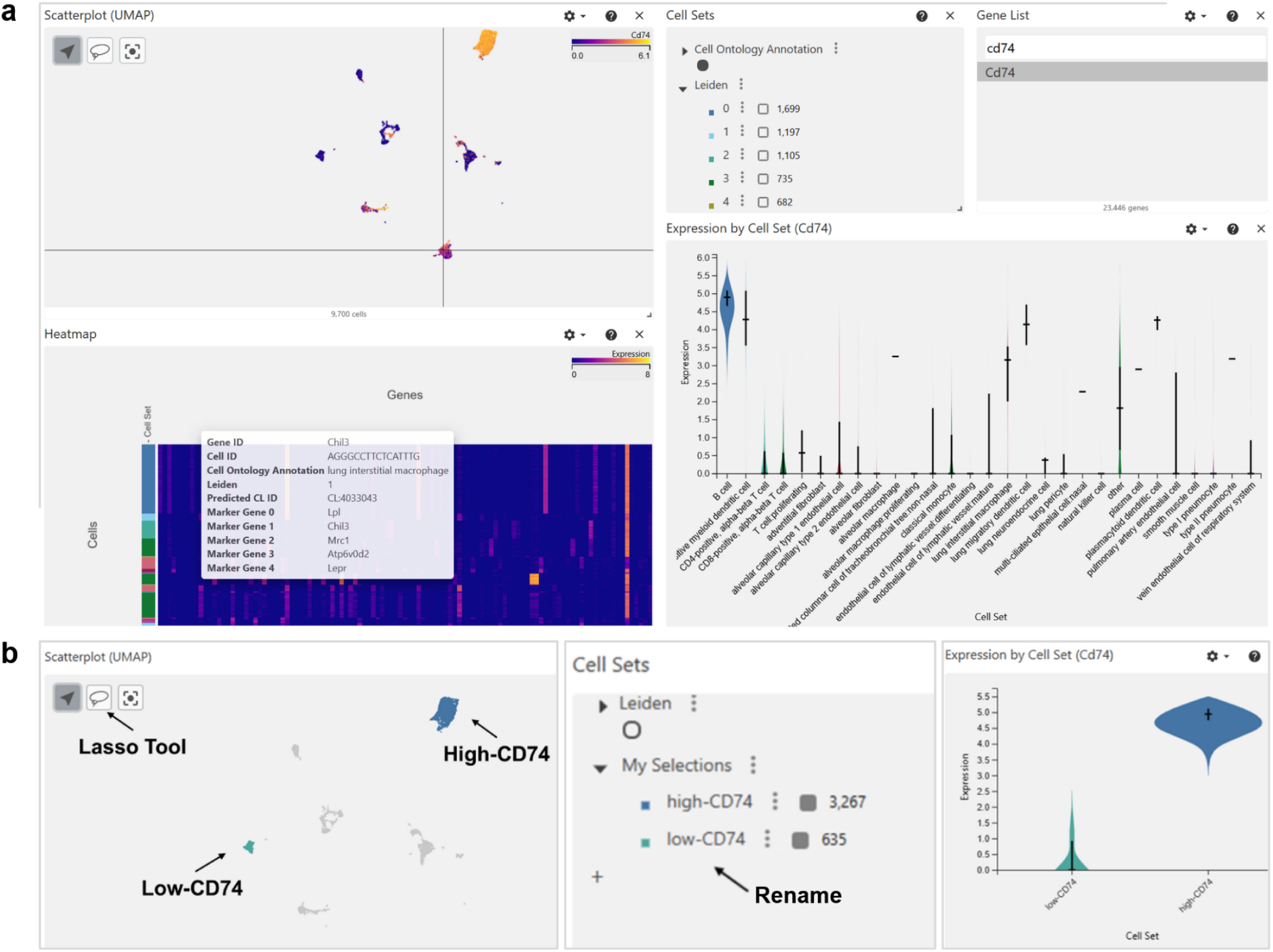
Visual analysis of gene expression in mouse lung tissue. **a.** The SenNet Portal supports interactive Vitessce visualizations of human and mouse datasets. Shown here are RNA-seq data from the left lung of a 5-month-old female C57BL/6J mouse. After searching for the candidate gene *Cd74*, the user can visualize its expression across all 9,700 cells in a UMAP plot (top left), heatmap (bottom), and by cell type in a violin plot (right). The Expression by Cell Set panel shows high *Cd74* expression in B cells (SenNet ID: SNT243.LTGW.532). **b.** The Vitessce lasso tool enables interaction with the UMAP plot and supports creating user-defined cell sets. The user can select cell regions that express Cd74-high (blue) and Cd74-low (teal). These new groups appear in the Cell Sets panel, where they can be managed (e.g., renamed). The violin plot visualizes the differential expression of *Cd74* between the user-defined sets. (SenNet ID: SNT243.LTGW.532).

#### Spatial Analysis

Understanding the link between pancreatic function and aging and disease is critical for senescence. Biomedical and computational scientists collaborating in an aging and senescence lab use the SenNet Portal to investigate this link further. From the Portal, they retrieve high resolution, multi-modal pancreas datasets, ranging from H&E histology to spatial transcriptomics (Xenium), and protein imaging (CODEX). Team members collaborate on an integrated workflow where pathologists use data to formulate questions about insulin production and computational researchers develop Jupyter notebooks that help answer them. In one workflow, they develop scripts that can overlay layers of data, perform cell segmentation, and apply spatial clustering to detect islets of Langerhans, the functional tissue units (FTUs) of the pancreas.^47^ The features of these islet are then analyzed, spatial abnormalities are explored, and healthy islets are separated from their senescent or dysfunctional counterparts. This workflow provides a generalizable framework that is useful for FTU detection in other organs through SenNet’s multi-layer datasets. Data and code developed as part of the Human Reference Atlas effort, including 2D illustrations of FTUs^47^, FTU segmentation algorithms^47,48^, and Cell Distance Explorer analyses^33^, support this effort and make it possible to explore SenNet data and results in the HRA.

#### Cost-efficient Profiling

Major barriers for researchers interested in lung aging include budgetary constraints and lack of access to extensive human tissue banks. The SenNet Portal provides these researchers with a free resource for studying how the lung’s cell type composition shifts across the lifespan. By querying the Portal’s APIs, researchers can retrieve all published single-nucleus RNA-sequencing datasets derived from human lung (**Fig. 5a**) and pass them through a custom processing pipeline, integrating the data using open-source Python modules, including Vitessce visualizations.^27,49–55^ They can then use annotation tools, like Azimuth^19^, CellTypist^56,57^, and other cell type annotation tools available via the HRA to classify cells across datasets (**Fig. 5b**). Data can further be grouped into age brackets to reveal shifts in the distribution of cell types across the lifespan, offering hints of potentially subclinical aging pathways at work before overt conditions emerge (**Fig. 5c**). Further extending the analysis, the researcher can control other donor factors, including BMI, blood type, medical/social history and compare the results with changes in cell type composition across other organs to determine whether age-related shifts are organ-specific or systemic. These resources offer an accessible and cost-effective way to track aging at both the cellular and cross-organ level without duplicating costly profiling efforts.

**Figure 5.**
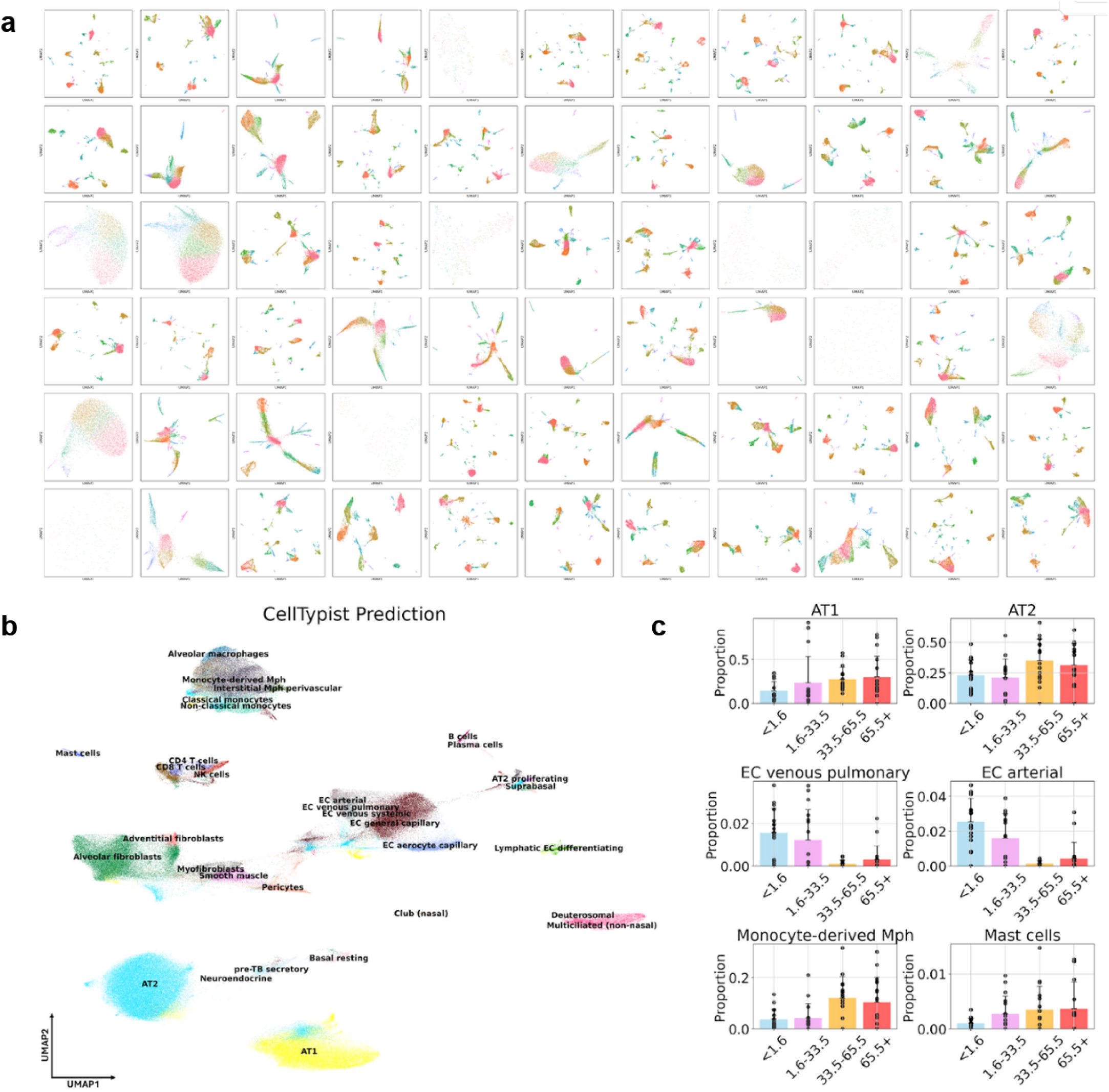
Integration reveals changes in human lung cell type composition across lifespan. **a.** UMAP embeddings for each of the 66 single-nucleus RNA-seq datasets may be visualized with software such as Vitessce within SenNet data portal functionality. **b.** Custom code is used to retrieve, filter, and integrate data across donors. Resulting harmonized CellTypist annotations are visualized on a single UMAP plot. (**c**) Donor metadata is used to group datasets by age quartile, demonstrating how proportions of putative epithelial, endothelial, and immune cells are influenced by donor factors such as age.

#### Cell Distance Distributions

Researchers interested in understanding immunology can use the HRA Cell Distance Explorer (CDE)^33^ to understand the location and distances between different cell types. **Fig. 6** shows one lymph node dataset with 1,826,800 segmented and annotated cells from Fan’s lab for a 25-year old female donor (SenNet Dataset ID: SNT584.CWSK.568). Data were obtained using Formalin-Fixed Paraffin-Embedded tissue samples cut from a lymph node in the stomach (**Fig. 6a**). A 49-plex marker panel optimized to identify B cells, T cells, macrophages, and other immune cell types was used to image the sample using PhenoCycler Fusion 1.0. The CDE was used to compute distances from all cell types to the nearest endothelial cells, which are then visualized as a violin plot showing distance distributions for each cell type (**Fig. 6b**). The CDE shows an interactive view of the data with cell-to-nearest-endothelial cell linkages allowing further inspection and closer examination of the data, see linkages in **Fig. 6c** that are colored by cell type. Examining the violin plot in the CDE shows the difference in absolute spatial location of different cell types to the endothelial cells, especially the different immune cells, i.e., the subtypes of B cells and T cells. The visualization also shows whether the spread of a cell type around the blood vessels is tighter or varied by looking at the interquartile ranges in the violin plots. The researcher can further examine changes in the distance distributions for samples from different donors, looking at changes in spatial cell organization with respect to certain anchor cells across age groups, sex, conditions, and regions.

**Figure 6.**
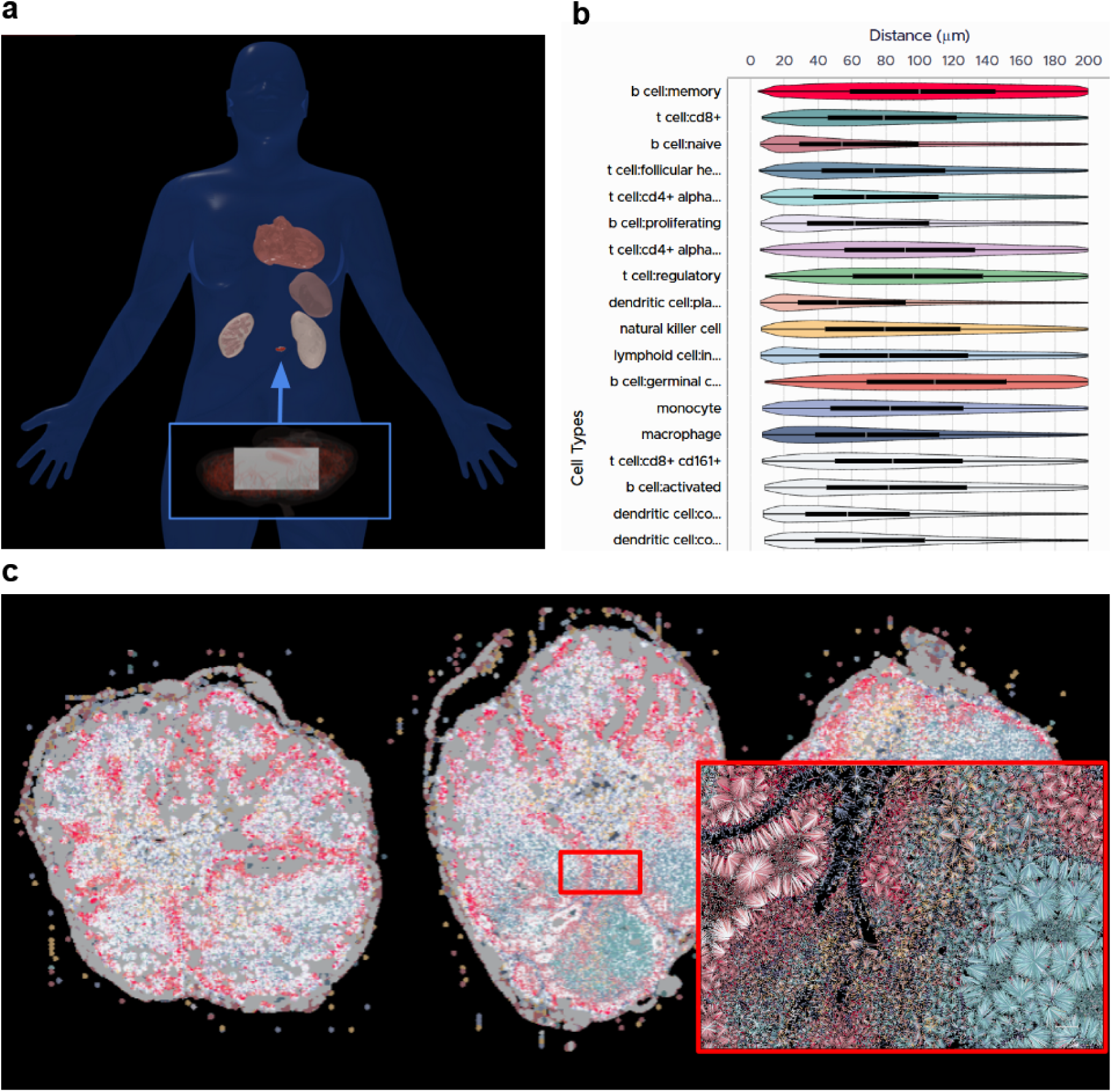
Cell distance distributions for immune cells in a lymph node of a 25 year old female donor. **a**. Lymph node tissue RUI-registered and shown in the context of the HRA female reference body. **b.** Violin plot of cell distance distributions for different color-coded cell types. **c.** Cell-to-nearest-endothelial cell distance visualization in the CDE interactive viewer.

Across donors spanning age, these distance distributions can reveal age-associated remodeling of vascular-associated lymph nodes. Spatial architecture of lymph node tissues in older donors are often described by a smaller node size, capsule thickening, fibrosis, lipid and hyaline deposition, and loss of clearly segregated B cell and T cell zones^58^. Furthermore, increased lymphatic vessel permeability and altered morphology of HEVs may further reshape perivascular localization patterns which results in altered immune cell entry. In the context of immune aging, such trends can be interpreted as spatial signatures of immunosenescence^59^. The changes in endothelial niches reshape where immune cells reside and impact antigen surveillance of functional microenvironments such as follicular and T-zone niches. In malignancies of lymph nodes, such as Angioimmunoblastic T cell lymphoma (AITL), there is significant remodeling of the normal lymph node architecture, characterized by infiltration of diverse immune cells, extensive follicular dendritic cell meshwork proliferation and prominent arborizing high endothelial venules (HEVs)^60^. Specifically, T follicular helper cells (Tfh) which are the tumor cell of origin are found in close proximity to the HEVs, highlighting deviations in cell distance distributions relative to healthy lymph nodes^61^.

There are a total of seven lymph node datasets that have been registered in the female reference body so they can be explored in the context of other organs in the SenNet EUI, go to https://data.sennetconsortium.org/ccf-eui and filter by female and Lymph Node. The CDE Python template was used to compute the cell-to-nearest-endothelial cell distance distributions.

### Utility and Usage

#### Expert Survey

Between April 8–15, 2026, we conducted an informal survey of experts in the biology of aging. The sample consisted of 3,169 researchers, drawn from the author lists and citation networks from three major publications in the field of senescence^3,14,15^. The survey attracted 190 respondents (144 complete). The respondents were predominantly PhD-holding experimentalists aged 35–64. 93% reported that senescence/aging is central or relevant to their work, indicating strong audience fit. However, 89% were outside the SenNet Consortium and most had low familiarity with the Portal, suggesting reach beyond the consortium to those with limited SenNet engagement.

Respondents were presented with a short version of the usage scenarios described above and asked to rank them in order of value. They placed a higher value on hypothesis generation and data quality benchmarking, followed by spatial analysis and lifespan profiling, with proximity analysis ranked last. Beyond these prompted scenarios, survey respondents suggested incorporating support for the following workflows: cross-species and cross-organ comparisons, upload-your-own-data scenarios, senotherapeutic target profiling, and extension into disease contexts such as early cancer, neurodegeneration, and diabetes.

The results of the survey could also be used to inform future development of the SenNet portal in four areas: data, workflows, user interface, and standardization. First, the most requested data gaps were higher quality spatial transcriptomics and proteomic assays, metabolomics/lipidomics, EHR linkage, and longitudinal age-matched donor cohorts with rich metadata. Genome-scale assays (WGS, long-read RNA-seq, 3D genomics, histone modification profiling, DNA damage/repair mapping) and integrated multi-omics were also frequently requested. Second, workflow-related requests included better spatial segmentation methods, integration of analysis software into the portal, and support for molecular interaction graphs. Third, respondents asked for improvements in the user-friendliness of the Portal’s Graphical User Interface (GUI) and searching and filtering capabilities. Finally, the respondents highlighted the importance of access to consistent data across papers and repeatedly called for robust and universal senescence biomarkers.

#### Usage Statistics

The SenNet-internal Data^62^ Ingest Board (**Fig. 7a**) lists the status of all 5,061 datasets (4,295 primary plus derivative datasets) that were uploaded whether or not they have been processed or published.

**Figure 7.**
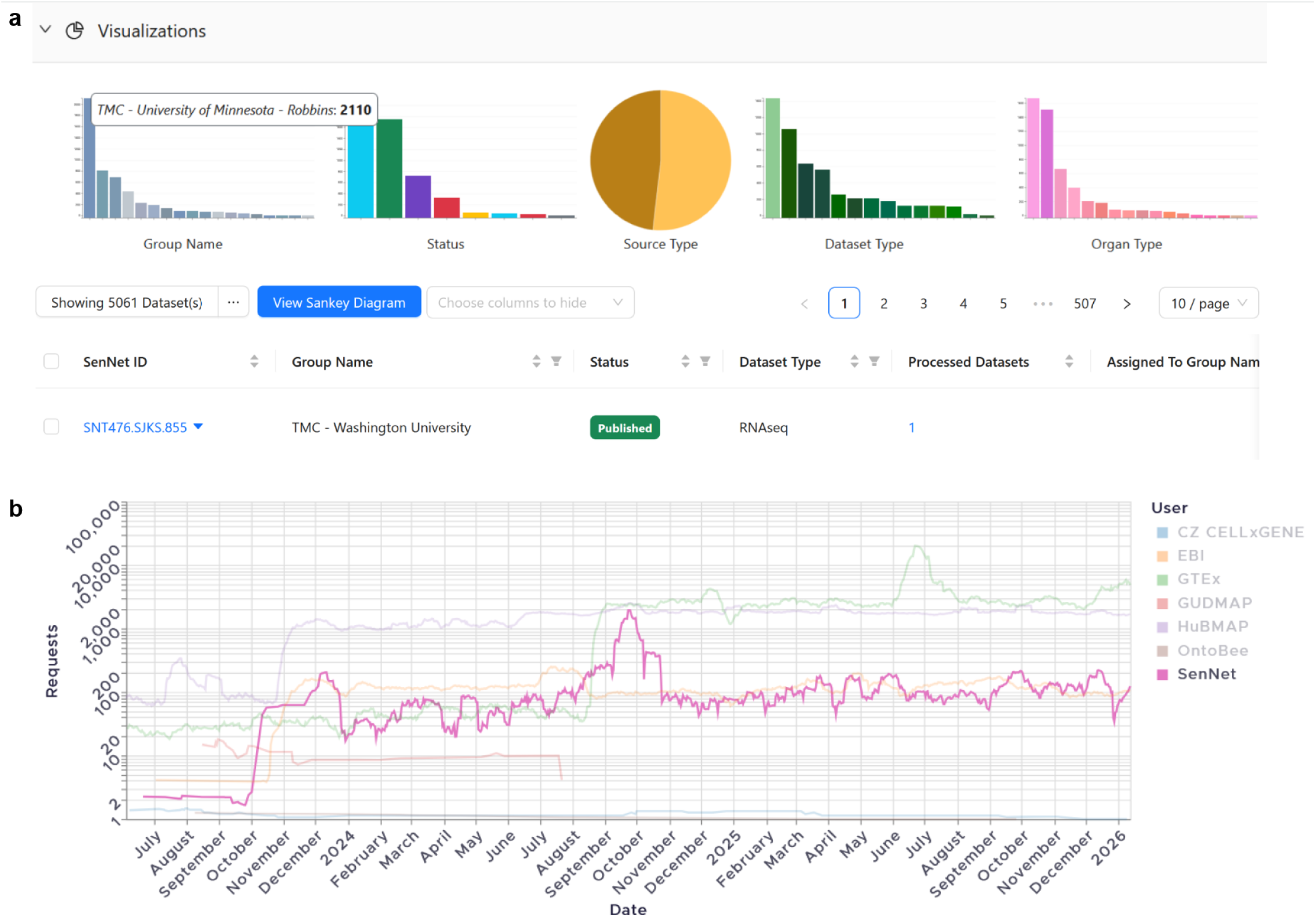
SenNet Dashboards. **a.** The internal Data Ingest Board lets SenNet members sort and filter datasets by authoring team, status, type, and 20+ other properties. **b.** The HRA Usage Dashboard shows HRA digital object requests from the SenNet Portal since June 2023, computed daily with a two week rolling average.

Usage for the SenNet Consortium public Web Portal (https://sennetconsortium.org), Data Portal (https://data.sennetconsortium.org), related sites and services (https://docs.sennetconsortium.org), is logged via AWS, Google, and GitHub. During the two year period of February 1, 2024 to February 1, 2026, from Google Analytics, the Web Portal has been viewed by 21,700 active users, the Data Portal viewed by 7,500 active users, and for the Documentation and Biomarkers site viewed by 3,200 active users, each increasing over time. Top referrers include: Google, Bing, trafficcheap.com, leadsgo.io, the-scientist.com, and nature.com. Most viewed pages include the base page (/), /search, /dataset, and /biomarkers with more than 5000 Data Portal users from the United States, and next more than 1,000 from China.

Also in the last two years, custom AWS logging of Downloads, APIs, and services reveals over 68.45 TiB of data has been downloaded, and more than 6,079,000 API calls have been made (including both direct use of published APIs via SDK/SmartAPI and those same calls that operate the portals). Top services include /search, /datasets. Top referrers are direct, auth.globus.org, and Google.

GitHub logged consortium contributions and usage for the infrastructure alone (not including individual site tools and algorithms) include over 40 repositories that have been viewed 4,600 times and cloned 5,300 times, see: https://github.com/x-atlas-consortia.

For the HRA Portal, between June 2023 and end of January 2026, 3,316,899 HRA Portal requests and 1,693,128 HRA API requests were fulfilled; the top-5 referrers are the GTEx Portal (https://gtexportal.org), the HuBMAP Entity API, the HuBMAP Data Portal, the SenNet Data Portal (https://data.sennetconsortium.org/search) (**Fig. 7b**), and EMBL-EBI (https://www.ebi.ac.uk). The 3D reference objects were accessed 34,704 times via the NIH3D website. The HRA OWL file was accessed 28,266 times via the NCBI BioPortal Ontology Browser (https://bioportal.bioontology.org/ontologies/HRA) and 39,577 times via the EBI OLS Ontology Browser (https://www.ebi.ac.uk/ols/ontologies/hra). 310 students registered for the VHMOOC and spent 5,652 hours reviewing materials, taking self-tests, and engaging in a community of practice.

In total, these reveal robust usage of information, data and services of the SenNet Consortium.

## Discussion

This Resource paper describes data, code, and tools that are of broad utility, interest, and significance to the senescence research community. Five usage scenarios exemplify how SenNet data and tools can be used to benchmark algorithms, generate hypotheses, perform spatial analyses, run cost efficient profiling, and explore senescent biomarkers in 3D. We conclude with a discussion of limitations and planned extensions.

In April 2026, 2,041 primary human and mouse datasets of the 3,247 total primary datasets for human and mouse were publicly available in the SenNet portal. The remaining datasets are either incomplete (e.g., in progress, uploading metadata, etc.) or are undergoing QA/QC protocols prior to publication. Experimental teams closely collaborate with the CODCC curators to finalize and publish these datasets via the SenNet Data Portal so they can be used by the worldwide scientific community.

We plan to make the usage scenarios presented here and others developed by SenNet members easy to access via the SenNet Data Portal and the HRA Portal. The usage scenarios help highlight unique data assets, provide hands-on guidance on how to use existing APIs and user interfaces, and showcase custom code developed by experimental teams.

The SenNet Portal will soon feature more “publication pages”. Publication pages highlight peer-reviewed publications and preprints that use consortium datasets.^63^ They make it easy to access and download at once, all datasets and code used in a paper. Two SenNet publication packages are planned in the *Cell Press* and *Nature* portfolio of journals. Papers in these packages are expected to have publication pages with a summary of publication-related information, a list of referenced SenNet datasets, plus relevant visualizations.

We use the SenNet Data Portal Usage data and dashboards for understanding and optimizing data and tool usage. Current dashboards (**Fig. 7a**) are only available to SenNet members, but a public dashboard showing the number of users and accesses over time is planned.

In parallel to the work presented here, the SenNet CODCC team is working on expanded spatial-omics releases and improved tools for senotype characterization by the consortium. This will enable assertional knowledge developed by the consortium to integrate fully in the same graph database as dataset metadata for further discovery and validation of senescence science.

## Acknowledgements

This research has been supported by the following awards:

The NIH Common Fund through the Office of Strategic Coordination/Office of the NIH Director:

- HuBMAP:

◦ OT2OD026671 and OT2OD033756 (K.B., A.B., Y.J., D.Q., B.W.H.)
◦ OT2OD026675 (P.D.B., J.C.S., B.H., B.S., M.S., D.B., G.R.S., A.B. as NIH JumpStart Award)
◦ OT2OD033759 (P.D.B., B.H., D.B., G.R.S., A.B. as NIH JumpStart Fellowship)
◦ 3OT2OD026682 (M.R.)
◦ 1OT2OD033761 (M.R.)
◦ OT2OD033760 (R.S.)
- SenNet:

◦ U24CA268108 (K.B., P.D.B., J.C.S., M.R., N.G., B.H., A.B., Y.J., D.Q., B.S., M.S., D.B., G.R.S., M.L.T., B.W.H.)
◦ U54AG076043 (A.E, R.F.; A.B. and Y.J. as supplement award)
◦ U54AG075936 (S.K., P.L.)
◦ U54AG079754 (J.B., S.T.P.)
◦ U54AG076041 (J.B., S.T.P.)
◦ U54AG075941 (S.D.)
- CFDE:

◦ OT2OD030545 (K.B., A.B., D.Q., B.W.H.)
◦ 1R03OD039970-01 (A.B., B.W.H., D.B., M.G.)

National Human Genome Research Institute [NHGRI):

- RM1HG011014 (R.S.)

National Institute of Diabetes and Digestive and Kidney Diseases:

- Kidney Precision Medicine Project: U01DK133090 (K.B., A.B., D.Q., B.W.H.)
- U2CDK114886 (K.B., A.B., Y.J., D.Q., B.W.H.)
- U24DK135157 (K.B., D.Q., B.W.H.)

National Institute of Mental Health (NIMH):

- RF1MH128876 (R.F.)
- RM1MH132648 (R.F.)

National Institute on Aging (NIA):

- U54AG079759 (R.F.)

National Cancer Institute (NCI):

- U01CA294514 (A.E., R.F.)
- UH3CA257393 (A.E., R.F.)
- U54CA274509 (R.F.)
- U54CA268083 (R.F.)
- R01CA245313 (R.F.)

K.B. is a co-director of and funded by the MacMillan Multiscale Human program by the Canadian Institute for Advanced Research (CIFAR). The funders had no role in study design, data collection and analysis, decision to publish or preparation of the manuscript. K.B. is also supported via a Stiftung Charité Visiting Fellowship via Berlin Institute of Health at Charité (BIH).

## Author Contributions

- K.B. led the writing of this paper and is a corresponding author with P.D.B., J.C.S., and B.W.H.
- D.Q. conducted user research with K.B. and wrote up results in the ***General User Needs***, ***Usage Scenarios***, and ***Usage: Expert Surveys*** sections, as well as ***Documentation and Demonstrations***.
- P.D.B. and J.C.S. led the development of the flexible hybrid cloud microservices architecture.
- J.C.S., P.D.B., B.S., M.S., D.B., and R.S. implemented key SenNet Consortium Portal and SenNet Data Portal functionality.
- K.B., B.W.H., A.B., Y.J., and D.Q. implemented key Human Reference Atlas Portal functionality.
- B.H. led and wrote up Data Submission and Ingestion.
- M.R. led the development of the data analysis pipelines.
- N.G. led the development of the Vitessce visualization tool.
- R.S. led the development of the Azimuth cell-type annotation tool.
- J.B. contributed the Data Quality Benchmarking usage scenario.
- S.-J.K and P.J.L. contributed the Hypothesis Generation usage scenario.
- S.D. contributed the Spatial Analysis usage scenario.
- S.P. contributed the Cost-efficient Profiling usage scenario.
- J.Y., A.E., N.F., R.F. contributed the Cell Distance Distributions usage scenario.
- K.B., P.D.B, J.C.S., D.Q., B.H., J.B., S.-J.K, S.D., S.P., A.B., Y.J., M.L.T., A.E., N.F., R.F., and B.W.H. wrote the paper. All other authors reviewed and commented on the paper.
- All SenNet Team authors have login credentials to the SenNet Data Portal on January 31, 2026.

## Competing Interests

### Core authors

In the past three years, R.S. has received compensation from Bristol-Myers Squibb, ImmunAI, Resolve Biosciences, Nanostring, 10X Genomics, Neptune Bio, and the NYC Pandemic Response Lab. R.S. is a co-founder and equity holder of Neptune Bio.

N.G. is a co-founder and equity owner of Datavisyn.

R.F. is scientific founder and adviser for IsoPlexis, Singleron Biotechnologies, and AtlasXomics. The interests of R.F. were reviewed and managed by Yale University Provost’s Office in accordance with the University’s conflict of interest policies.

All other authors declare no competing interests.

## Full list of SenNet Team authors

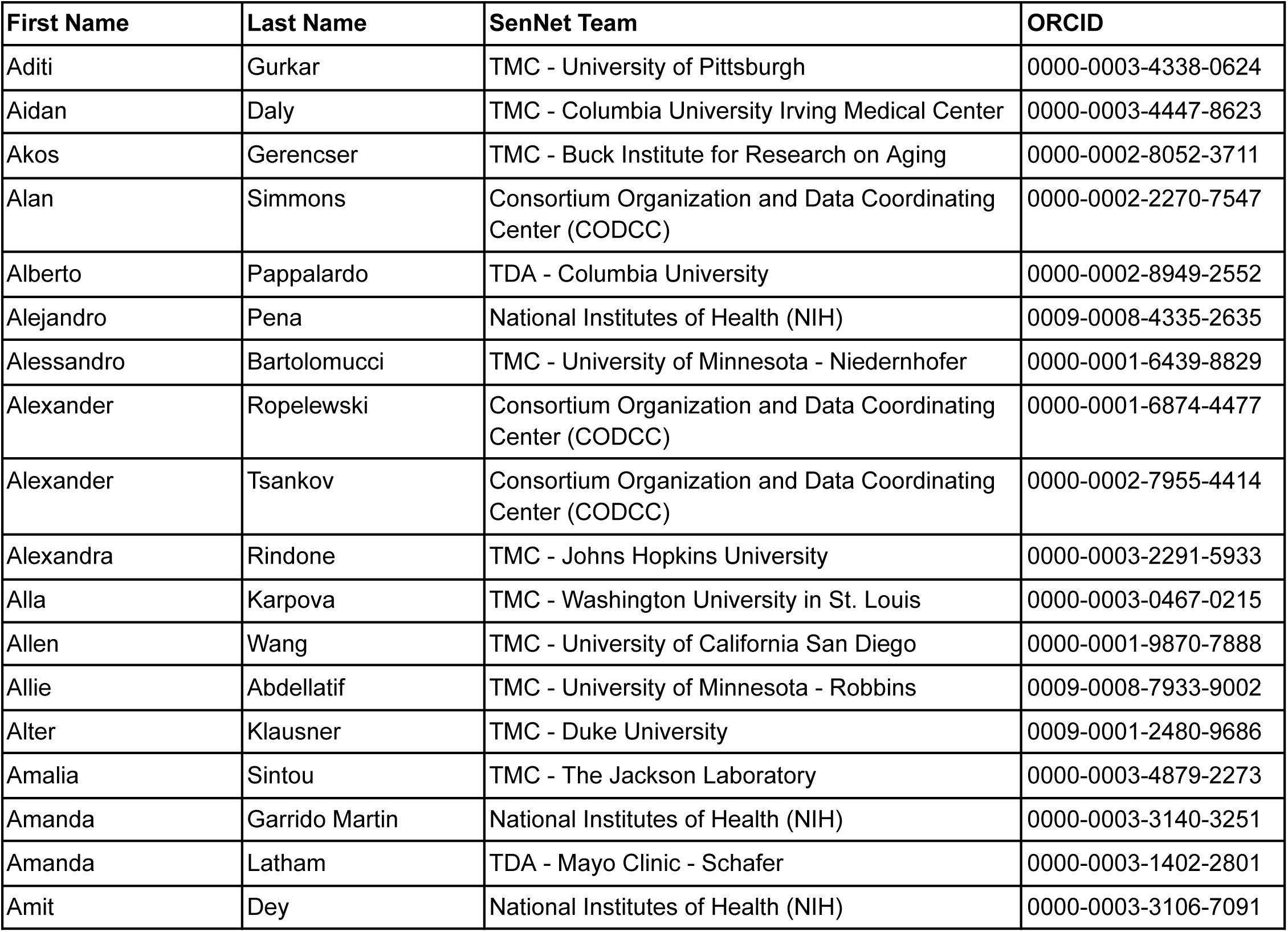
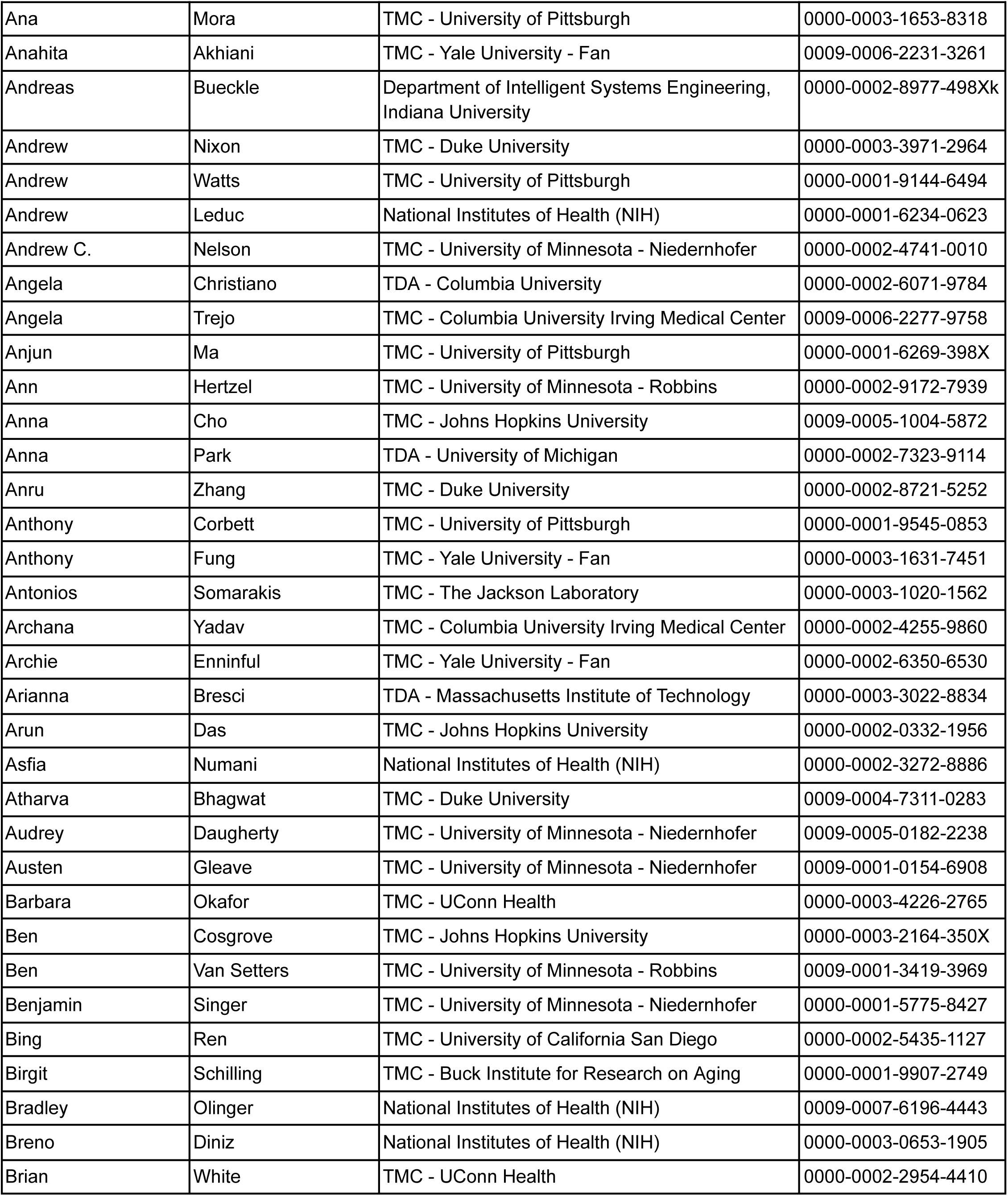
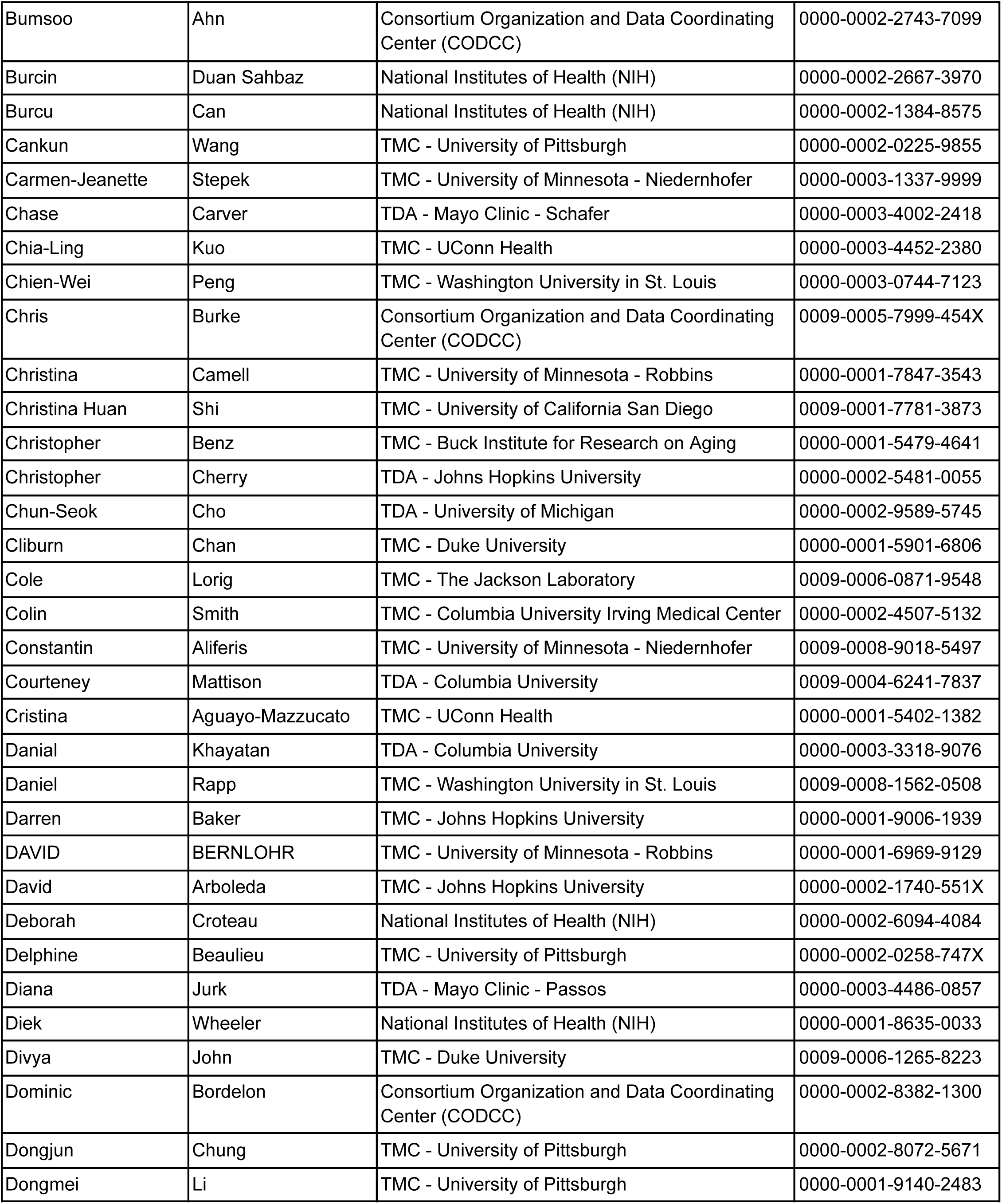
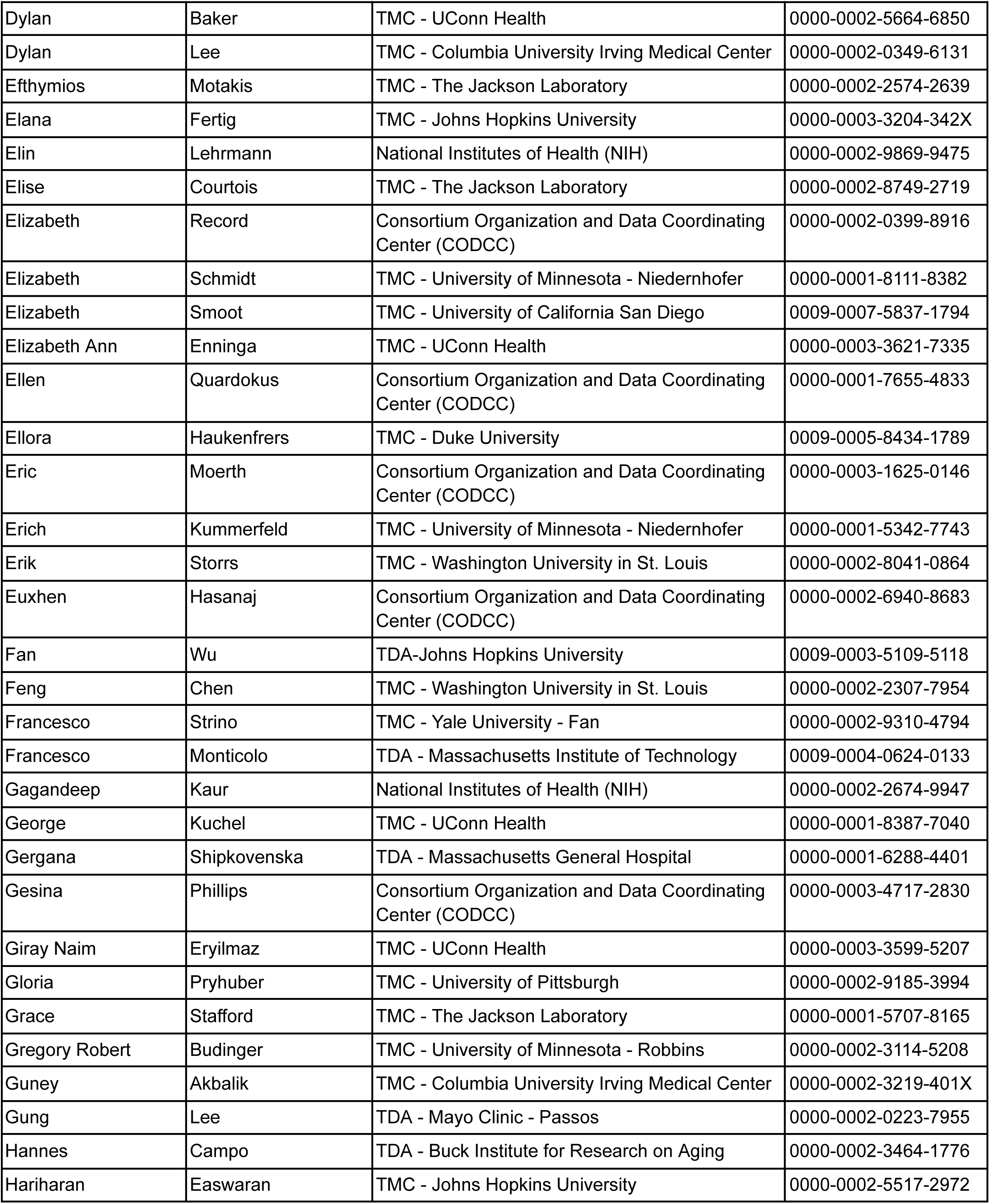
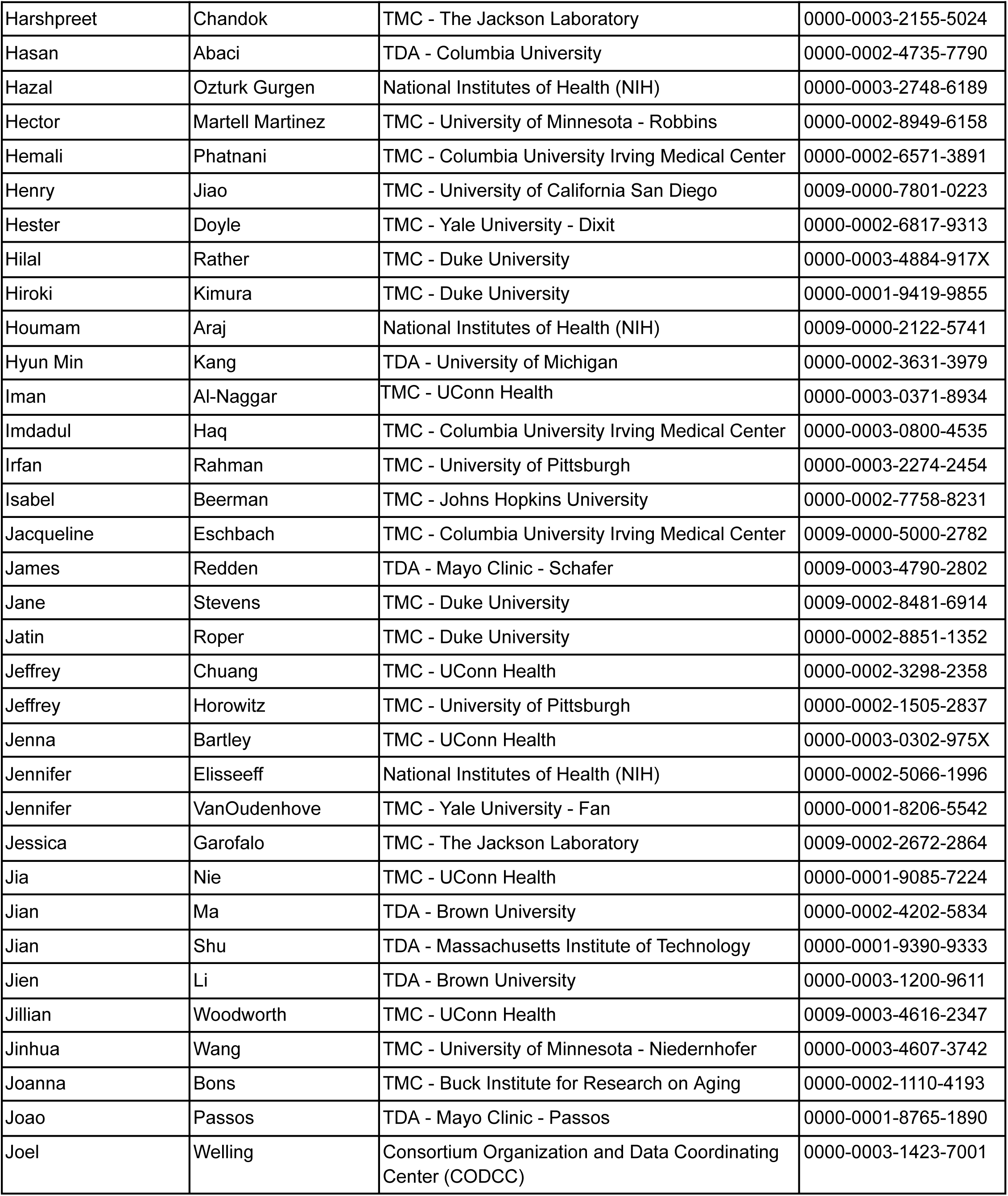
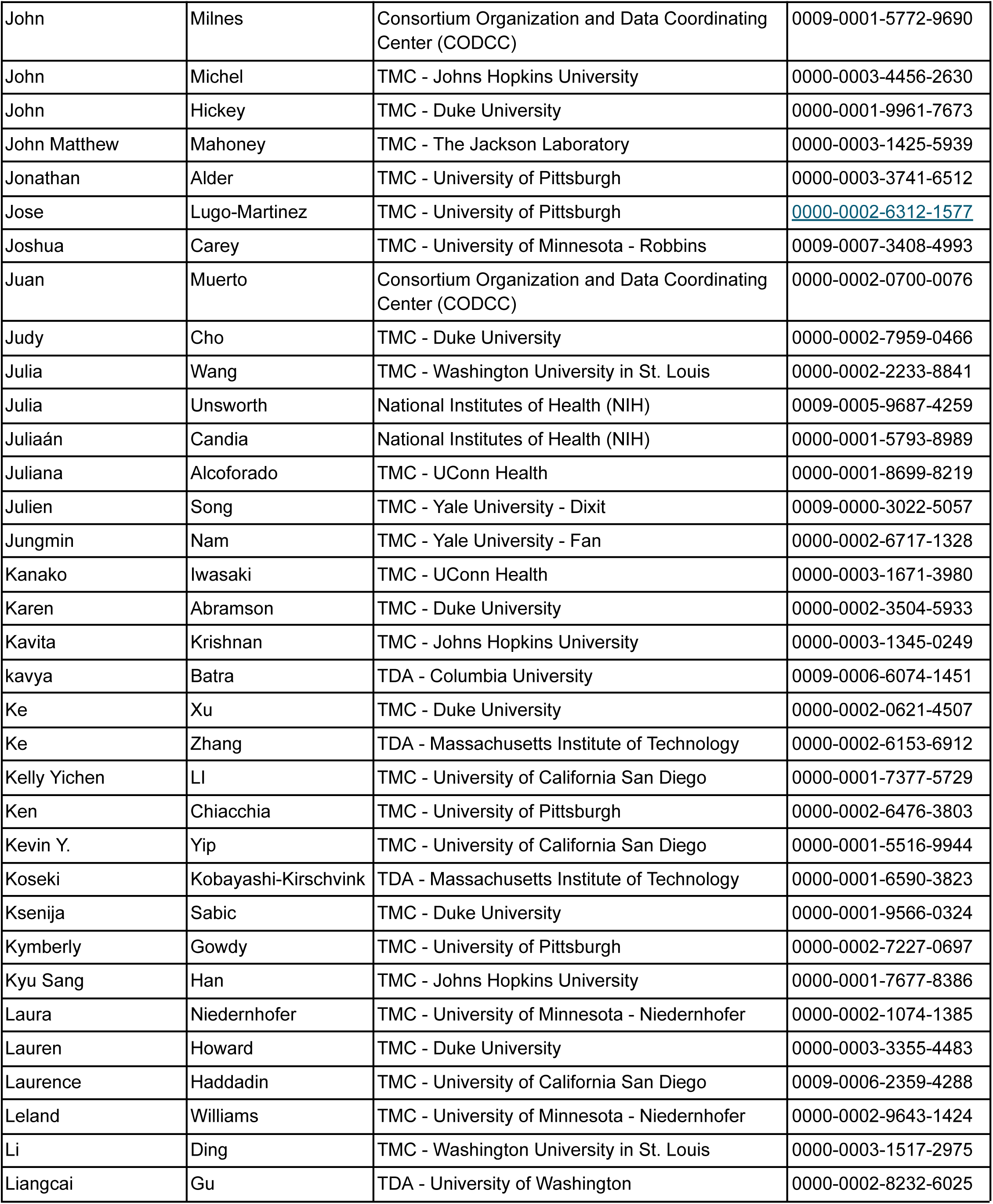
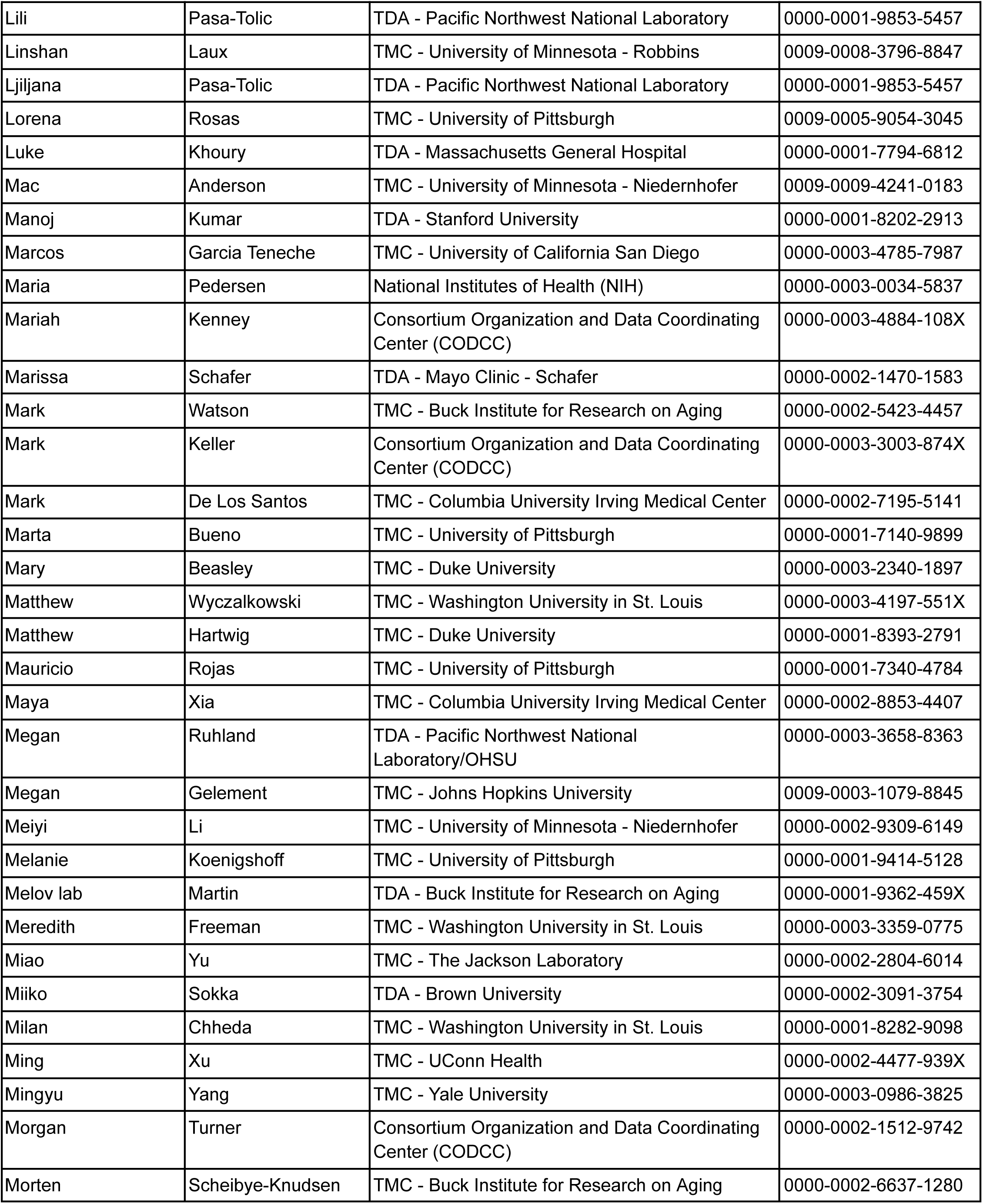
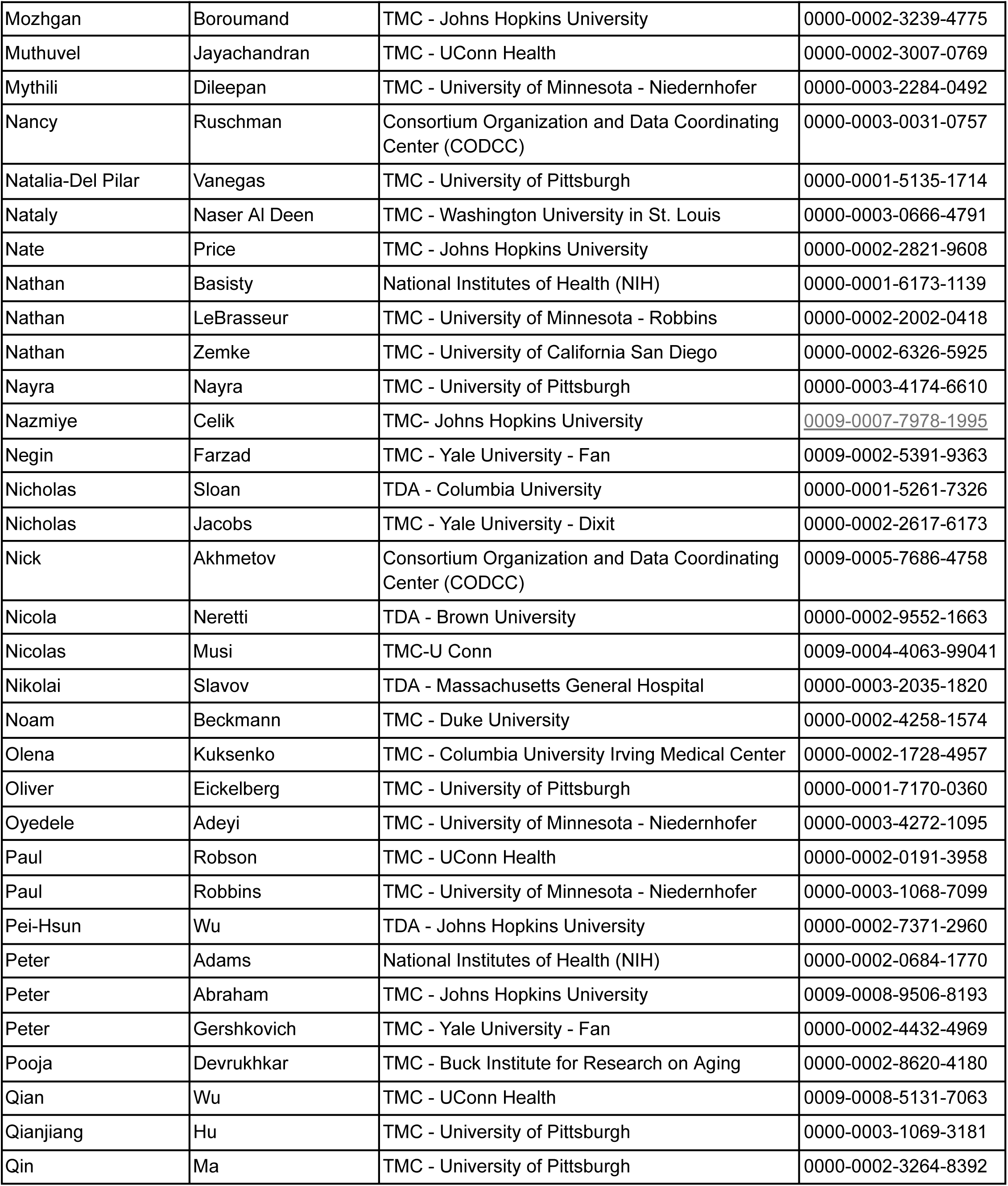
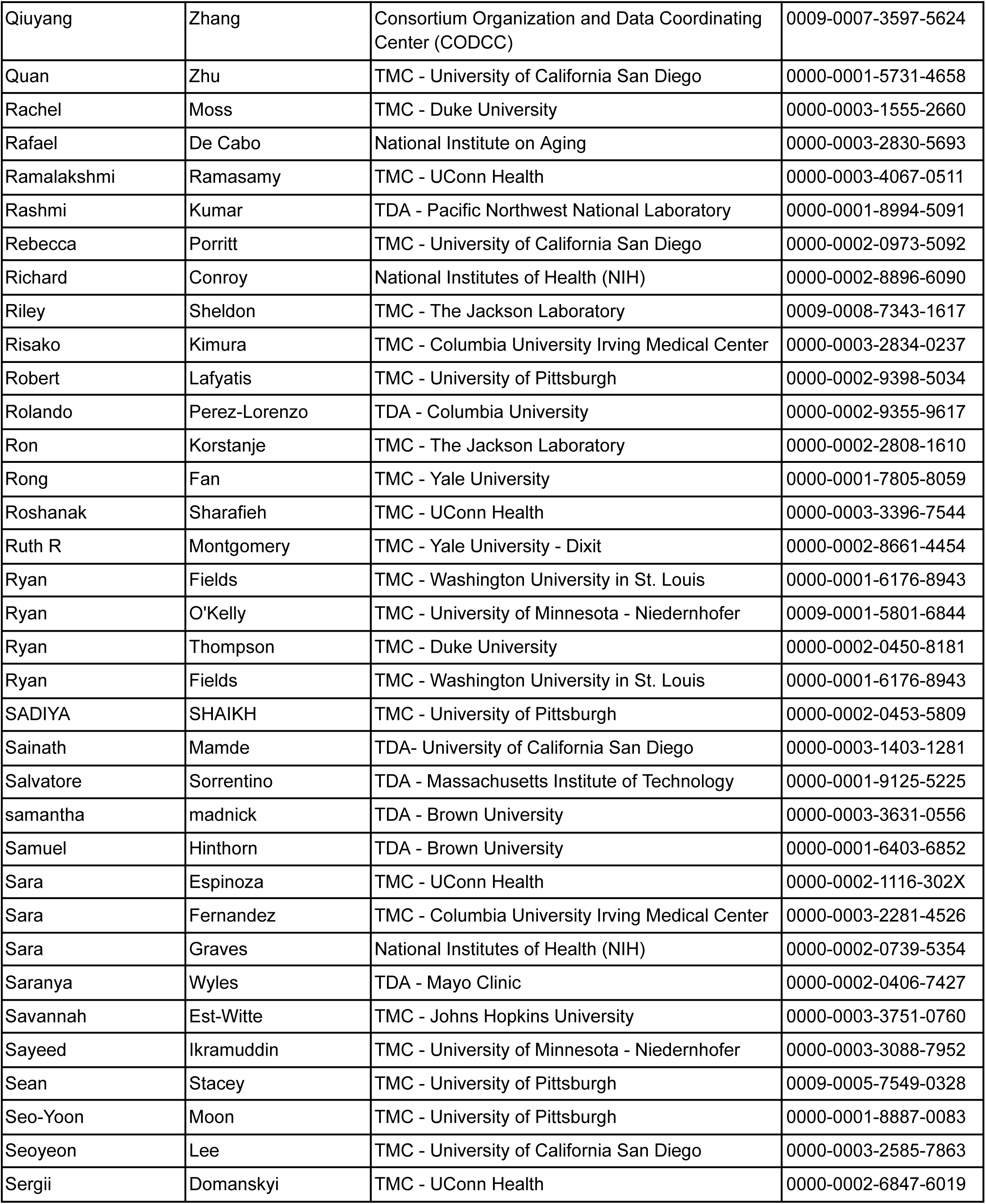
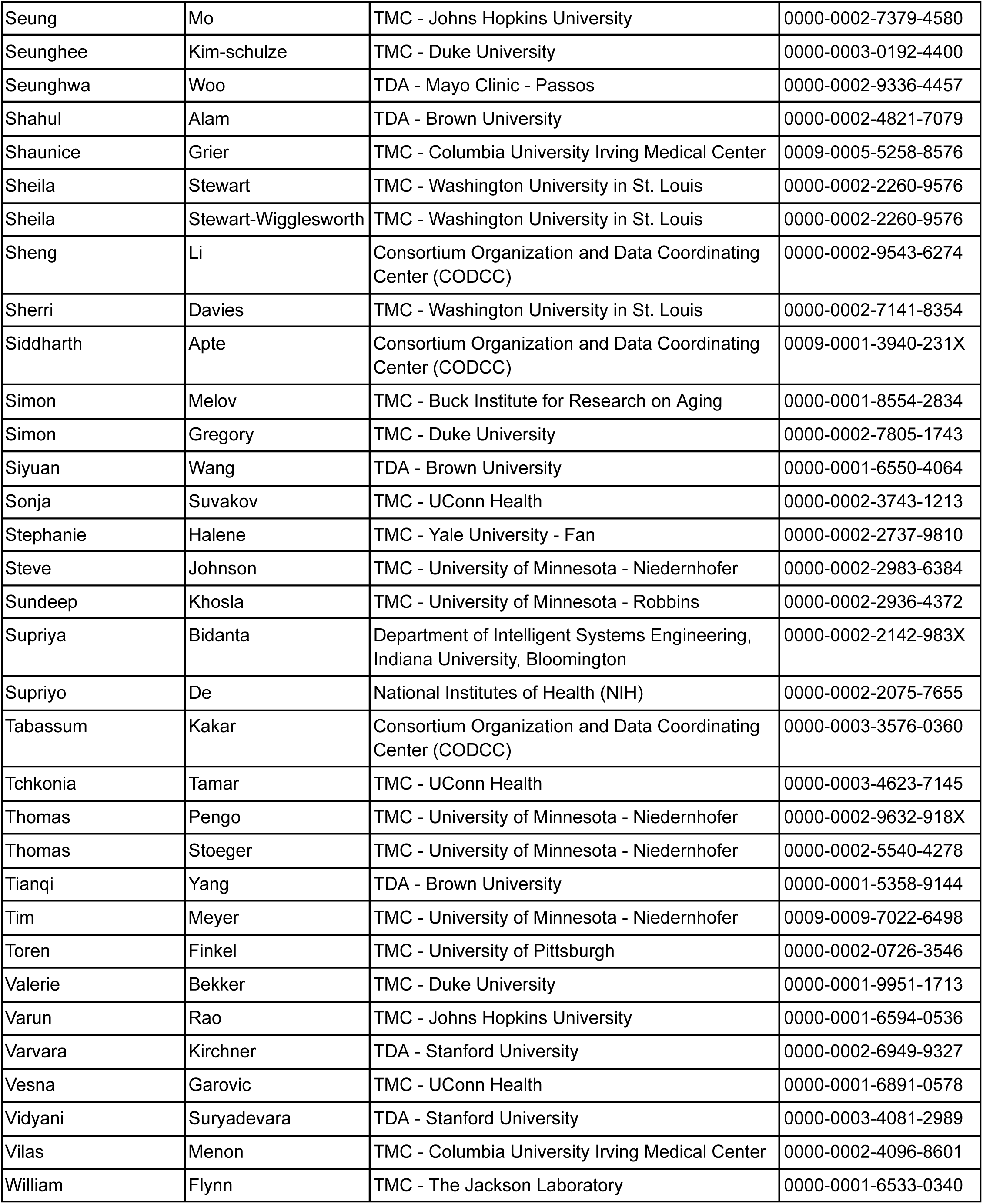
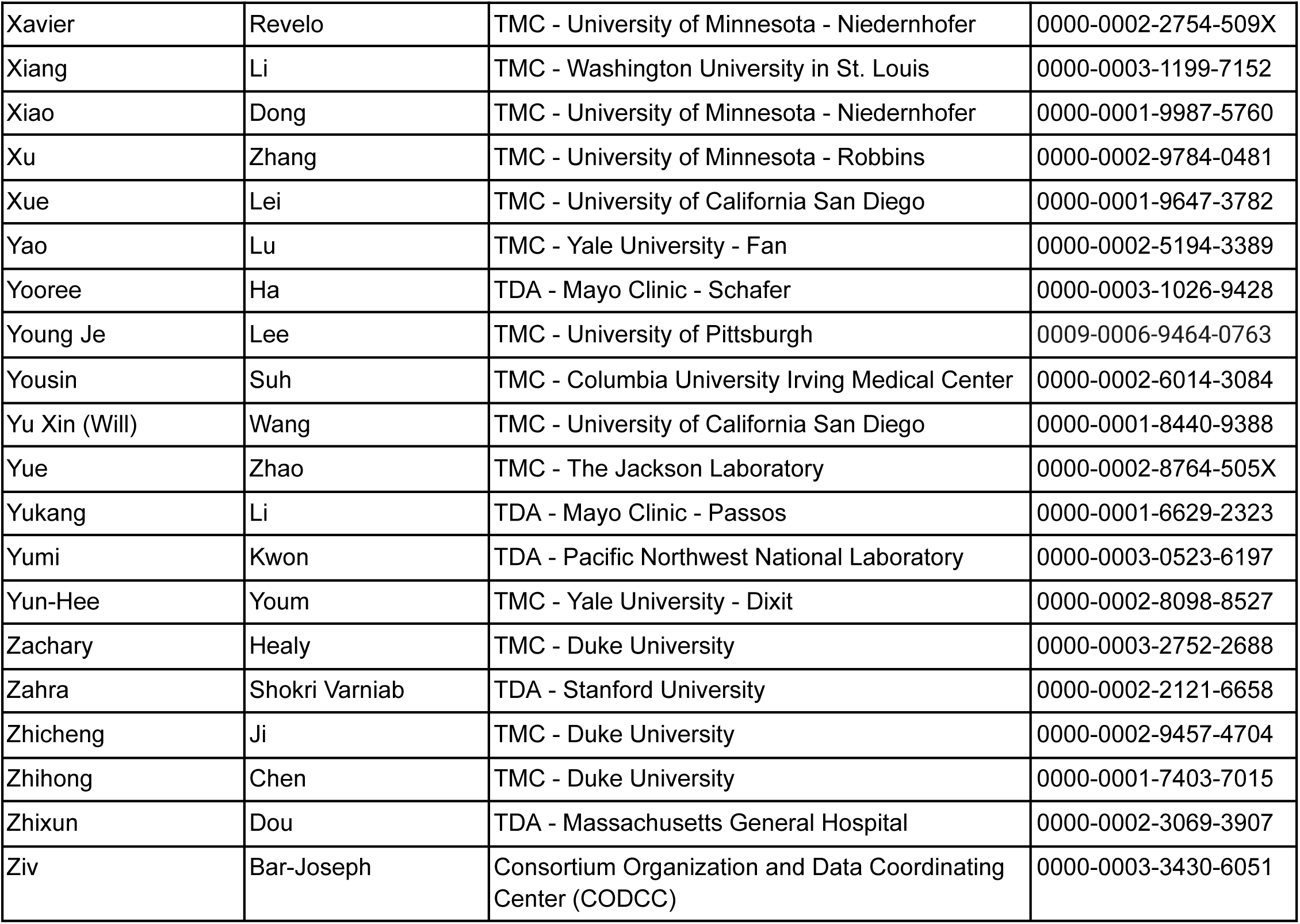

## Methods

### Data Quality Benchmarking (Josh Bartz)

A single-cell RNA-seq (scRNA-seq) computational pipeline was developed for processing and benchmarking the datasets. The pipeline processed raw scRNA-seq datasets through a 7-step workflow: Cleaning, Integration, Clustering, Cell Type Identification, Calculating Scallop scores, Calculating Global Coordination Level (GCL), and Visualization. The workflow allowed comparison of different methods of measuring transcriptional noise. The following datasets were used by this pipeline to generate **Fig. 3**: SNT246.PGHH.774, SNT449.GRWG.598, and SNT397.SPGW.436. Seurat was used for data cleaning, integration, clustering, and cell type identification (steps 1-4). Since GCL calculations can be computationally demanding, the overall GCL analysis was split into smaller chunks which was then run on high performance computers (HPCs) using SLURM job scheduler. The complete code (predominantly in the R programming language) for all seven steps is available via GitHub (se*e **Code Availability***).

### Spatial Analysis (Sergii Domanskyi)

A set of 12 Jupyter notebooks demonstrating data access, analysis, and visualization workflows for the SenNet Data Portal were created (see ***Code Availability***). These notebooks showcase practical applications of spatial analysis across computational biology, with a focus on tissue architecture, cell-cell interactions, and disease-related cellular changes for high-resolution, multi-modal datasets including spatial transcriptomics, imaging, and single-cell RNA sequencing data from diverse human and mouse tissues. The demos also provide accessible entry points into senescence research for discovering datasets, performing gene expression analysis, and integrating multi-modal imaging data. The raw and processed datasets used throughout these notebooks were retrieved from the SenNet Portal. These include datasets SNT576.PXPP.452, SNT484.VLRN.777, SNT443.KFNS.239, SNT574.MMQC.683, SNT594.JCFN.856, and SNT227.HLMG.672 from collection SNT793.SZRS.468, dataset SNT544.XHGB.538 from collection SNT947.NXPB.793, and datasets SNT268.LXPG.784, SNT324.BDTT.263, and SNT469.JJLS.674 from collection SNT566.NMTV.379.

### Cost-efficient Profiling (Sam Peters)

Relevant single-cell datasets were queried using the parameterized search endpoint of the SenNet Search API, and expression matrix files of qualifying datasets were downloaded using the Entity and Assets APIs. Datasets were then processed and integrated with Python (v3.9.23) using scanpy (v1.10.3)^42^ and anndata (v0.10.9)^40^. Basic quality control metrics across cells and genes were calculated. Then, basic thresholds were set to filter out noisy data: cells that had fewer than 500 transcript counts; cells with more than 10% mitochondrial content; and genes expressed in fewer than 3 cells across samples. Data were normalized and log1p transformed prior to scaling for dimensionality reduction. Highly variable genes were annotated in a batch-wise manner, followed by scaling and PCA. Data were then integrated to adjust the principal components (PC) using Harmony (v0.0.10)^46^, and 30 integrated PCs were used to calculate 30 nearest neighbors. Finally, a UMAP^64^ embedding was computed for visualization and CellTypist (v1.7.1)^47,48^ was applied to annotate cell types based on the Human Lung Cell Atlas (HLCA, v1.0) model. The processed data were used to examine the effect of age on cell type composition. All code is publicly available (see ***Code Availability***). Processed data from following SenNet datasets were used in this usage scenario example: SNT246.PGHH.774, SNT397.SPGW.436, SNT449.GRWG.598, SNT498.XKRG.893, SNT233.XMTD.574, SNT722.MQMW.937, SNT673.HKBD.656, SNT759.KMJC.586, SNT627.TGCJ.324, SNT985.MXCW.829, SNT266.DBBP.698, SNT493.RSKL.732, SNT776.QQGW.623, SNT536.TGBM.294, SNT967.KBJF.497, SNT867.PMJD.853, SNT478.QGSL.332, SNT394.ZPJN.579, SNT826.HWLM.338, SNT792.KVKQ.344, SNT372.NMWW.264, SNT577.KHFG.572, SNT367.FWNH.537, SNT328.LPZQ.663, SNT657.STXR.948, SNT593.SCSG.422, SNT299.DVNG.578, SNT933.RZWH.988, SNT339.DQDB.424, SNT362.BRTK.755, SNT647.WHJN.292, SNT489.CCVQ.324, SNT829.BMNR.662, SNT794.NKKN.985, SNT669.GCNV.685, SNT864.KHJR.253, SNT479.PCMQ.829, SNT253.MKNV.455, SNT236.VDFB.848, SNT654.XSRN.726, SNT467.NJWW.933, SNT676.SXTP.348, SNT984.MSST.526, SNT773.DRXG.427, SNT945.WCZH.229, SNT497.HRKZ.873, SNT644.VGFZ.956, SNT977.VVTW.775, SNT468.XMQP.379, SNT658.LQWB.482, SNT872.FZFV.997, SNT432.BQVB.669, SNT767.MZKB.693, SNT627.QHCP.732, SNT959.SGTC.264, SNT625.CWGD.777, SNT289.MKGJ.729, SNT648.KPTH.546, SNT983.FMRG.925, SNT587.CNQX.768, SNT835.GVZP.228, SNT757.BGFN.798, SNT387.XHRC.584, SNT473.ZJKW.727, SNT887.TMFM.723, SNT493.TKJD.678.

### Cell Distance Distributions

Using the Cell Distance Explorer python template at https://portal.hubmapconsortium.org/templates/hra_cde_tutorial, a Python Jupyter notebook was created that loads the cell table containing x-y coordinates and cell type assignments for a lymph node dataset of a 25-year-old female donor (SenNet Dataset ID for LN00560 sample: SNT584.CWSK.568) and visualizes the cell distance distributions for 1,826,800 cells. The cell table for this sample was processed in prior work^33^ and was used for this analysis and demo. All code is publicly available (see ***Code Availability***) and the sample cell table used can be downloaded from the CDE Spatial Omics Gallery (https://apps.humanatlas.io/cde/gallery/lymphnode-codex-yale). For a non-code user, the same analysis can also be computed and visualized using the browser-based application available at https://apps.humanatlas.io/cde.

## Data Availability

All SenNet data are available via the SenNet Portal (https://portal.hubmapconsortium.org). Azimuth references can be accessed at https://azimuth.hubmapconsortium.org. HRA data and code are available at the HRA Portal (https://humanatlas.io).

## Code Availability

Code is available on three different GitHub organizations: (1) https://github.com/sennetconsortium is the primary SenNet organization, (2) https://github.com/cns-iu is the organization owned by the Cyberinfrastructure for Network Science Center at Indiana University, and initial experimental HRA code starts here, (3) https://github.com/x-atlas-consortia was created to host cross-consortia code, including hra-kg, hra-pop, hra-apps, and hra-api. APIs, available as RESTful web services, support data ingest, querying, and delivery of metadata, see details at https://docs.sennetconsortium.org/apis. Code from the ***Usage Scenarios*** section is available at https://github.com/joshbartz55/SenNetSingleCellRNAseqDemo (Data Quality Benchmarking), https://github.com/TheJacksonLaboratory/sennet-demo (Spatial Analysis), https://github.com/TheJacksonLaboratory/sennet-demo/blob/main/notebooks/demo-snrna-seq-all-human-lung.ipynb (Cost-Efficient Profiling), and https://github.com/x-atlas-consortia/hra-notebooks/blob/main/notebooks/hra-cde-sennet.ipynb (Cell Distance Distributions).

## References

1. Roy, A. L. et al. A Blueprint for Characterizing Senescence. Cell 183, 1143–1146 (2020).

2. Li, S. et al. Advancing biological understanding of cellular senescence with computational multiomics. Nat. Genet. 57, 2381–2394 (2025).

3. Suryadevara, V. et al. SenNet recommendations for detecting senescent cells in different tissues. Nat. Rev. Mol. Cell Biol. 25, 1001–1023 (2024).

4. Jain, S. et al. Advances and prospects for the Human BioMolecular Atlas Program (HuBMAP). Nat. Cell Biol. 25, 1089–1100 (2023).

5. HuBMAP Consortium. The human body at cellular resolution: the NIH Human Biomolecular Atlas Program. Nature 574, 187–192 (2019).

6. Roy, A. L. et al. Accelerating a paradigm shift: The Common Fund Single Cell Analysis Program. Sci. Adv. 4, eaat8573 (2018).

7. Turner, M. L., et al. HuBMAP Data Portal: A Resource for Multi-Modal Spatial and Single-Cell Data of Healthy Human Tissues. Preprint at 10.48550/arXiv.2511.05708 (2025).

8. Wilkinson, M. D. et al. The FAIR Guiding Principles for scientific data management and stewardship. Sci. Data 3, 160018 (2016).

9. Honick, B. Getting Started with Data Submission. SenNet Consortium https://docs.sennetconsortium.org/libraries/ingest-validation-tools/upload-guidelines/getting-started/ (2024).

10. Record, E. Human Reference Atlas Standard Operating Procedures Glossary. Zenodo 10.5281/zenodo.17664992 (2025) doi:10.5281/zenodo.17664992.

11. Börner, K. et al. Tissue registration and exploration user interfaces in support of a human reference atlas. *Commun*. Biol. 5, 1369 (2022).

12. SenNet Consortium. SenNet Provenance Graph. SenNet https://docs.sennetconsortium.org/provenance/.

13. Herr II, B. W. et al. Specimen, biological structure, and spatial ontologies in support of a Human Reference Atlas. Sci. Data 10, 171 (2023).

14. He, S. & Sharpless, N. E. Senescence in Health and Disease. Cell 169, 1000–1011 (2017).

15. SenNet Consortium et al. NIH SenNet Consortium to map senescent cells throughout the human lifespan to understand physiological health. *Nat*. Aging 2, 1090–1100 (2022).

16. Börner, K. et al. Human BioMolecular Atlas Program (HuBMAP): 3D Human Reference Atlas construction and usage. Nat. Methods 1–16 (2025) doi:10.1038/s41592-024-02563-5.

17. Bueckle, A. & Qaurooni Fard, D. Using the Standalone Registration User Interface. 10.5281/zenodo.14346543 (2024) doi:10.5281/zenodo.14346543.

18. Bueckle, A. & Qaurooni Fard, D. Using the Embedded Registration User Interface. 10.5281/zenodo.14346342 (2024) doi:10.5281/zenodo.14346342.

19. Hao, Y. et al. Integrated analysis of multimodal single-cell data. Cell 184, 3573–3587.e29 (2021).

20. Keller, M. S. et al. Vitessce: integrative visualization of multimodal and spatially resolved single-cell data. Nat. Methods 22, 63–67 (2025).

21. HuBMAP Consortium. HuBMAP Data Submission Metadata. HuBMAP https://docs.hubmapconsortium.org/metadata (2026).

22. HuBMAP Consortium. Source Murine. SenNet https://docs.sennetconsortium.org/registration/schemas/source-murine/ (2026).

23. SenNet Consortium. Cellular Senescence Network (SenNet) Method Development Community - research workspace on protocols.io. protocols.io https://www.protocols.io/workspaces/cellular-senescence-network-sennet-method-develo (2026).

24. New York Genome Center Mapping Component. Azimuth: App for reference-based single-cell analysis. https://azimuth.hubmapconsortium.org (2026).

25. Cyberinfrastructure for Network Science Center. Human Reference Atlas (HRA) Portal 3D Multiscale Biomolecular Human Reference Atlas Construction, Visualization, and Usage. https://humanatlas.io/ (2026).

26. Patro, R., Duggal, G., Love, M. I., Irizarry, R. A. & Kingsford, C. Salmon provides fast and bias-aware quantification of transcript expression. Nat. Methods 14, 417–419 (2017).

27. Wolf, F. A., Angerer, P. & Theis, F. J. SCANPY: large-scale single-cell gene expression data analysis. Genome Biol. 19, 15 (2018).

28. Granja, J. M. et al. ArchR is a scalable software package for integrative single-cell chromatin accessibility analysis. Nat. Genet. 53, 403–411 (2021).

29. Chen, H. & Murphy, R. F. Evaluation of cell segmentation methods without reference segmentations. Mol. Biol. Cell 34, ar50 (2023).

30. Zhang, T., Chen, H., Lee, Y. J., Ruffalo, M. & Murphy, R. F. SPRM: Spatial process and relationship modeling for multiplexed images. Bioinforma. Adv. vbag019 (2026) doi:10.1093/bioadv/vbag019.

31. Qu, Y. et al. Single-cell and spatial detection of senescent cells using DeepScence. Cell Genomics 5, 101035 (2025).

32. Bueckle, A. et al. Cell Type Populations for 3D Anatomical Structures of the Human Reference Atlas. Sci. Data 10.1038/s41597-026-06642-4 (2026) doi:10.1038/s41597-026-06642-4.

33. Jain, Y. et al. Exploring endothelial cell environments across organs in spatially resolved omics data. 2025.09.23.678129 Preprint at 10.1101/2025.09.23.678129 (2025).

34. Manz, T. et al. Viv: multiscale visualization of high-resolution multiplexed bioimaging data on the web. Nat. Methods 19, 515–516 (2022).

35. Börner, K. et al. Anatomical structures, cell types and biomarkers of the Human Reference Atlas. Nat. Cell Biol. 23, 1117–1128 (2021).

36. Quardokus, E. M. et al. Organ Mapping Antibody Panels: a community resource for standardized multiplexed tissue imaging. Nat. Methods 20, 1174–1178 (2023).

37. SenNet Consortium. Section 4 - RUI Tissue Blocks. SenNet https://docs.sennetconsortium.org/data-submission/Section4.html (2026).

38. Bueckle, A. HRA Organ Gallery on Meta Store. Meta Quest Store https://www.meta.com/en-gb/experiences/hra-organ-gallery/5696814507101529/ (2025).

39. SenNet Consortium. Google Scholar Page for SenNet Consortium. https://scholar.google.com/citations?user=wvo05TQAAAAJ (2026).

40. Bartz, J., Jung, H., Wasiluk, K., Zhang, L. & Dong, X. Progress in Discovering Transcriptional Noise in Aging. Int. J. Mol. Sci. 24, 3701 (2023).

41. Amit, G., Vaknin Ben Porath, D., Levy, O., Hamdi, O. & Bashan, A. Global coordination level in single-cell transcriptomic data. Sci. Rep. 12, 7547 (2022).

42. Levy, O. et al. Age-related loss of gene-to-gene transcriptional coordination among single cells. Nat. Metab. 2, 1305–1315 (2020).

43. Ibañez-Solé, O., Ascensión, A. M., Araúzo-Bravo, M. J. & Izeta, A. Lack of evidence for increased transcriptional noise in aged tissues. eLife 11, e80380 (2022).

44. Wang, J. et al. A transcriptome-based human universal senescence index (hUSI) robustly predicts cellular senescence under various conditions. Nat. Aging 5, 1159–1175 (2025).

45. Jochems, F. et al. The Cancer SENESCopedia: A delineation of cancer cell senescence. Cell Rep. 36, 109441 (2021).

46. Tao, W., Yu, Z. & Han, J.-D. J. Single-cell senescence identification reveals senescence heterogeneity, trajectory, and modulators. Cell Metab. 36, 1126–1143.e5 (2024).

47. Bidanta, S. et al. Functional tissue units in the Human Reference Atlas. Nat. Commun. 16, 1526 (2025).

48. Cyberinfrastructure for Network Science Center. Human Reference Atlas Portal: Functional Tissue Unit (FTU) Illustrations. https://humanatlas.io/2d-ftu-illustrations (2025).

49. McKinney, W. Data Structures for Statistical Computing in Python. SciPy 2010 10.25080/Majora-92bf1922-00a (2010) doi:10.25080/Majora-92bf1922-00a.

50. Virshup, I., Rybakov, S., Theis, F. J., Angerer, P. & Wolf, F. A. anndata: Access and store annotated data matrices. J. Open Source Softw. 9, 4371 (2024).

51. Virshup, I. et al. The scverse project provides a computational ecosystem for single-cell omics data analysis. Nat. Biotechnol. 41, 604–606 (2023).

52. Hunter, J. D. Matplotlib: A 2D Graphics Environment. Comput. Sci. Eng. 9, 90–95 (2007).

53. Harris, C. R. et al. Array programming with NumPy. Nature 585, 357–362 (2020).

54. Waskom, M. L. seaborn: statistical data visualization. J. Open Source Softw. 6, 3021 (2021).

55. Korsunsky, I. et al. Fast, sensitive and accurate integration of single-cell data with Harmony. Nat. Methods 16, 1289–1296 (2019).

56. Domínguez Conde, C., et al. Cross-tissue immune cell analysis reveals tissue-specific features in humans. Science 376, eabl5197 (2022).

57. Xu, C. et al. Automatic cell-type harmonization and integration across Human Cell Atlas datasets. Cell 186, 5876–5891.e20 (2023).

58. Cakala-Jakimowicz, M., Kolodziej-Wojnar, P. & Puzianowska-Kuznicka, M. Aging-Related Cellular, Structural and Functional Changes in the Lymph Nodes: A Significant Component of Immunosenescence? An Overview. Cells 10, 3148 (2021).

59. Liu, Z. et al. Immunosenescence: molecular mechanisms and diseases. Signal Transduct. Target. Ther. 8, 200 (2023).

60. Chiba, S. & Sakata-Yanagimoto, M. Advances in understanding of angioimmunoblastic T-cell lymphoma. Leukemia 34, 2592–2606 (2020).

61. Zhu, M. et al. Single-cell transcriptomic and spatial analysis reveal the immunosuppressive microenvironment in relapsed/refractory angioimmunoblastic T-cell lymphoma. Blood Cancer J. 14, 218 (2024).

62. SenNet Consortium. SenNet Data Ingest Board. https://ingest.board.sennetconsortium.org/ (2026).

63. HuBMAP Consortium. Publications | HuBMAP Peer-Reviewed Research and Preprints. https://portal.hubmapconsortium.org/publications (2026).

64. McInnes, L., Healy, J., Saul, N. & Großberger, L. UMAP: Uniform Manifold Approximation and Projection. J. Open Source Softw. 3, 861 (2018).

